# Deep conservation of cis-regulatory elements and chromatin organization in echinoderms uncover ancestral regulatory features of animal genomes

**DOI:** 10.1101/2024.11.30.626178

**Authors:** Marta S. Magri, Danila Voronov, Saoirse Foley, Pedro Manuel Martínez-García, Martin Franke, Gregory A. Cary, José M. Santos-Pereira, Claudia Cuomo, Manuel Fernández-Moreno, Alejandro Gil-Galvez, Rafael D. Acemel, Periklis Paganos, Carolyn Ku, Jovana Ranđelović, Maria Lorenza Rusciano, Panos N. Firbas, José Luis Gómez-Skarmeta, Veronica F. Hinman, Maria Ina Arnone, Ignacio Maeso

**Author notes:** Posthumous author. These authors have contributed equally to this work (Marta S. Magri, Danila Voronov, Saoirse Foley).

## Abstract

Despite the growing abundance of sequenced animal genomes, we only have detailed knowledge of regulatory organization for a handful of lineages, particularly flies and vertebrates. These two groups of taxa show contrasting trends in the molecular mechanisms of 3D chromatin organization and long-term evolutionary dynamics of cis-regulatory element (CREs) conservation. To help us identify shared versus derived features that could be responsible for the evolution of these different regulatory architectures in animals, we studied the evolution and organization of the regulatory genome of echinoderms, a lineage whose phylogenetic position and relatively slow molecular evolution has proven particularly useful for evolutionary studies. First, using PacBio and HiC data, we generated new reference genome assemblies for two species belonging to two different echinoderm classes: the purple sea urchin *Strongylocentrotus purpuratus* and the bat sea star *Patiria miniata*. Second, we characterized their 3D chromatin architecture, identifying TAD-like domains in echinoderms that, like in flies, do not seem to be associated with CTCF motif orientation. Third, we systematically profiled CREs during sea star and sea urchin development using ATAC-seq, comparing their regulatory logic and dynamics over multiple developmental stages. Finally, we investigated sea urchin and sea star CRE evolution across multiple evolutionary distances and timescales, from closely related species to other echinoderm classes and deuterostome lineages. This showed the presence of several thousand elements conserved for hundreds of millions of years, revealing a vertebrate-like pattern of CRE evolution that probably constitutes an ancestral property of the regulatory evolution of animals.

## Introduction

Animals are arguably the most morphologically disparate eukaryotic lineage. The development of this diversity of body plans rests on a common principle: a highly complex spatio-temporal regulation of gene expression. Accordingly, in contrast to their unicellular relatives ^1^, gene expression in animals often occurs through the coordinated action of multiple cis-regulatory elements (CREs), which can be located at very long distances from their target genes. However, in different animal groups, this tight regulation in time and space can be achieved through very different strategies, which in turn are reflected in profound differences in the way their genomes are organized from a regulatory perspective ^2–8^. How have these diverse gene regulation strategies evolved in animals? Which ones are ancestral and which ones are novel? Recent works using comparative functional genomics have started to delve into these problems ^6,7,9–12^, but the number of lineages sampled is still quite limited. Although different lines of evidence support that long-range regulation is probably ancestral to animals ^13–17^, there are marked differences between lineages in terms of the architectural proteins involved, so the ancestral molecular mechanisms underlying the evolution of distal regulatory interactions remain unclear. In vertebrates, distant regions of the genome can be brought together via loop extrusion, a mechanism in which cohesin and the zinc-finger protein CTCF play fundamental roles. Chromatin loops are anchored by convergently oriented CTCF binding sites, and this particular orientation of the CTCF-bound motifs is crucial for the formation of these 3D chromatin structures ^18–21^. However, in flies, interactions between areas of the genome with similar chromatin states and additional lineage-specific architectural proteins such as Cp190, BEAF-32 and Su(Hw) have a much larger impact on 3D contacts than CTCF ^22^. Furthermore, CTCF function in flies is independent of the orientation of its cognate CTCF motifs ^23^. We also have a very limited understanding of CRE conservation at deep evolutionary distances. A large fraction of CREs are either evolutionarily short-lived or subjected to high rates of turnover ^24–29^ and in lineages such as flies and nematodes CRE sequences are only conserved over short evolutionary timescales ^26,30–33^. In stark contrast with these rapidly changing CREs, we also know that in other lineages, certain enhancers can be staggeringly old, with a few examples that predate the split between different bilaterian animal phyla and even between cnidarians and bilaterians, dating back to more than 600 mya ^34–36^. The biological and evolutionary significance of this extreme sequence conservation ^37,38^ is a particularly challenging question because, so far, the only animal lineage with widespread deeply conserved CREs are the vertebrates, with several thousands of them maintained across jawed vertebrate species for over 450 million years ^39–43^. The extent of deep CRE conservation in many lineages other than flies, nematodes and vertebrates remains unexplored ^6,8,9,11,12^ or it is impossible to assess because they are phylogenetically isolated taxa ^7,44^.

Echinoderms are particularly well suited to fill this gap and provide an ideal framework for macroevolutionary comparisons of their regulatory genome. Their extant species span a wide range of phylogenetic distances and evolutionary timescales and their genomes have retained several ancestral features, such as a high conservation of developmental gene families ^45^ and conserved microsyntenic associations ^13,16,46^. Therefore, echinoderms play a pivotal role in our understanding of the genomes of deuterostome and bilaterian animal ancestors.

In the last decades, multiple works have compared developmental gene regulatory networks (GRNs), which require knowledge of genes, their products and the CRE controlling these genes, between different echinoderm lineages, especially between sea star and sea urchin species, revealing a remarkable conservation of regulatory nodes, but also considerable rearrangement of network architecture after 500 Mya of evolution^47^. In addition, classical small-scale gene-centered studies ^48,49^ and more recently, through genome-wide comparisons ^50^ have also investigated CRE evolution between relatively closely related sea urchin species. However, very little is known about genome-wide trends of CRE evolution between deeply diverged echinoderm lineages and between echinoderms and other deuterostomes, while the 3D regulatory organization of their genomes remains unexplored.

Here, we performed a comprehensive study of the bat sea star and the purple sea urchin regulatory genomes and 3D chromatin organization and assessed their evolution across major echinoderm lineages to investigate how the functional genome architecture of different animal lineages evolved.

## Results

### New Bat Sea Star and Purple Sea Urchin Reference Genome Assemblies & Gene Annotations

To investigate gene regulatory evolution in echinoderms and comprehensively profile CREs together with 3D chromatin organization, we first generated new reference genome assemblies for two of the best developmentally characterized echinoderm species, the bat sea star, *Patiria miniata*, a representative of the class Asteroidea, and the purple sea urchin, *Strongylocentrotus purpuratus*, from the class Echinoidea (Fig. 1a). We sequenced the samples using PacBio Sequel, achieving 123x coverage for *S. purpuratus*, and 125x for *P. miniata* and generated HiC data for scaffolding. These new *P. miniata* v3.0 (GCA_015706575.1) and *S. purpuratus* v5.0 (GCA_000002235.4) genome assemblies have a total size of 608 Mb and 921 Mb respectively (Fig. 1b) and 96% and 96.3% BUSCO completeness (Extended Data Fig. 1a). The scaffold N50 was of 23 Mb for *P. miniata* and 37 Mb for *S. purpuratus*, which represents a 437.26 and 88.89 fold improvement in contiguity respectively from previously available genome assemblies (Extended Data Fig. 1b,c). 90% of the assemblies were contained within the largest 23 *P. miniata* and 22 *S. purpuratus* scaffolds (i.e. scaffold L90 values).

**Figure 1:**
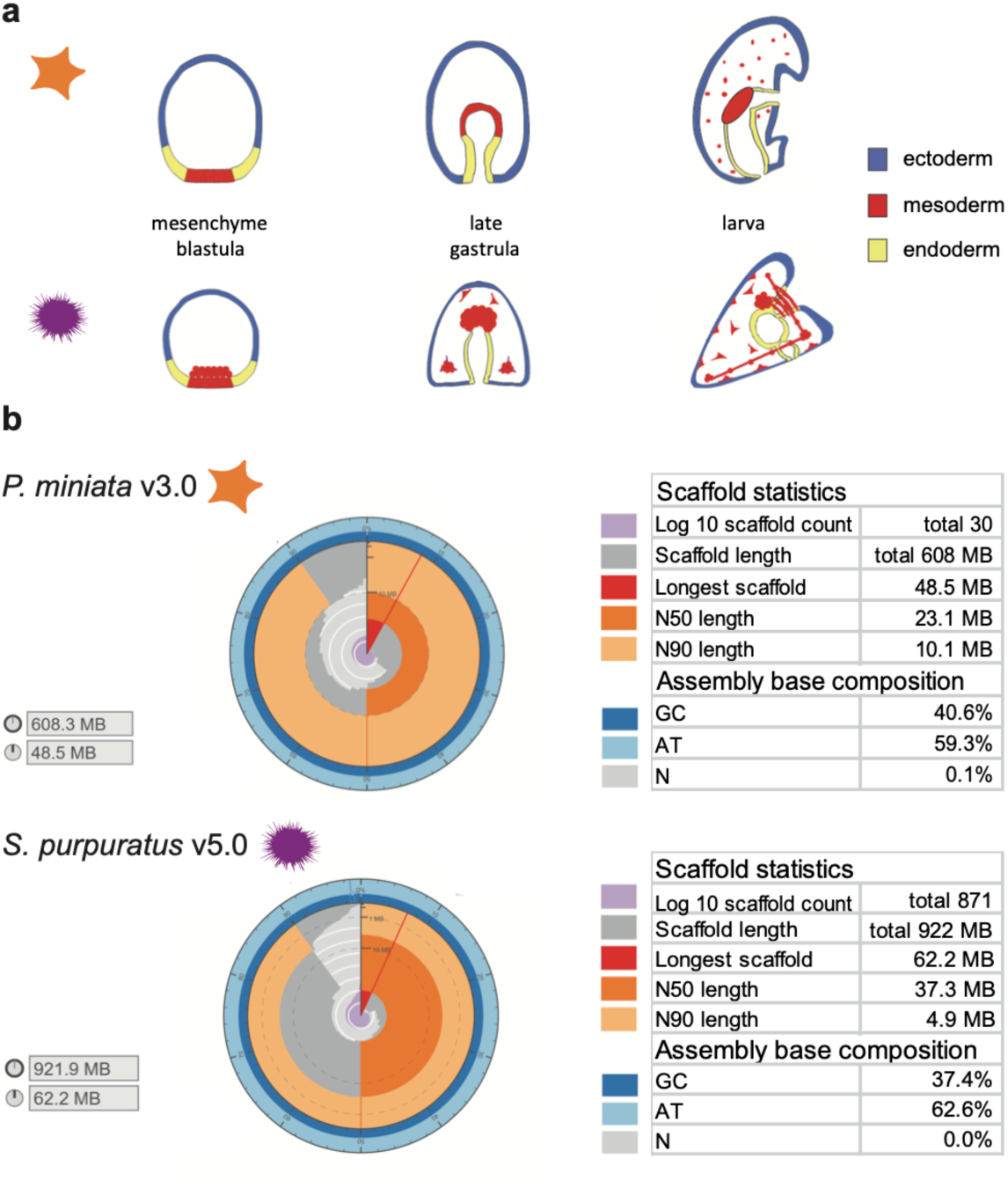
New Reference Genome Assemblies. **a**, Schematic drawings showing the germ layers and their derivatives of the equivalent developmental stages chosen for the generation of ATAC-seq (all stages) and Hi-C (late gastrula stage) data in the bat sea star (*P. miniata*) and purple sea urchin (*S. purpuratus*). **b,** Snailplots of general metrics (scaffold total length, scaffold count, longest scaffold, N50 and N90 values, and base compositions) of the new assemblies of the bat sea star (GCA_015706575.1) and purple sea urchin (GCA_000002235.4).

The genome annotations, both obtained using the NCBI eukaryotic genome annotation pipeline ^51^, contain 33,503 gene models (of which 27,447 are protein coding), with 38,426 mRNA transcripts for *S. purpuratus* and 27,818 gene models (of which 23,044 are protein coding), with 35,403 mRNA transcripts for *P. miniata*.

### Chromatin folding in Echinoderms

To characterize chromatin folding in our two echinoderm species, we used HiC data from late gastrula stage of sea star and sea urchin embryos and compared them with data from other animal lineages. We first observed that at the sub-megabase scale the *P. miniata* and *S. purpuratus* genomes fold in self-insulated domains reminiscent of Topologically Associated Domains (TADs) observed in other animals (Fig. 2a,b). We used insulation scores and boundary scores to define a comprehensive set of putative TADs and TAD boundaries, identifying 2,108 and 3,009 boundaries in each species, respectively. Importantly, previously identified conserved microsyntenic pairs, which are usually associated with developmental gene families, such as Pax1/9, Foxa, Tbx2/3, and Egr ^13,16^, that also show a high abundance of conserved non-coding sequences ^3^, were included within these TADs, supporting that these domains could have a role in long-range regulation in echinoderms (Extended Data Fig. 2, see also next section on CRE conservation). We then investigated which molecular mechanisms could be responsible for boundary formation in echinoderms. First, we confirmed that both *P. miniata* and *S. purpuratus* genomes have orthologs of the two main proteins involved in loop extrusion and boundaries in vertebrates, CTCF and cohesin subunit SA1-3 (Extended Data Fig.3a,b). In the case of CTCF, the orthologs of both echinoderms and their sister lineage, the hemichordates, were more divergent than those of other bilaterian phyla, and were characterized by the loss of one (in hemichordates) or two (in echinoderms) of the eleven Zn-finger domains ancestrally present in bilaterian CTCF proteins (Extended Data Fig.3c), as it had been previously noticed in another sea urchin species, *Hemicentrotus pulcherrimus* ^52^. Next, we checked if CTCF could be playing a similar function to vertebrates in echinoderm TAD boundaries, by looking for CTCF motif enrichment in *P. miniata*, *S. purpuratus* and a vertebrate species as a positive control (the zebrafish *Danio rerio*). Both sea star and sea urchin showed a CTCF motif enrichment in boundaries, although much lower than in the vertebrate case, especially in the case of *S. purpuratus* where CTCFs motifs were absent in nearly half of the boundaries (Fig. 2c,d). We next assessed if these CTCF motifs in echinoderm TAD boundaries displayed the typical vertebrate pattern with divergently oriented CTCF binding sites at each side of the boundaries. In stark contrast with zebrafish, the orientation of sea star and sea urchin CTCF motifs at boundaries did not show any orientation pattern and their distribution was indistinguishable from the controls (Fig. 2e). Consistent with this, there was no enrichment of CTCF motifs at sea star and sea urchin loop anchors, suggesting that CTCF plays a different role in these species than in vertebrates (Extended Data Fig 4a,b).

**Figure 2:**
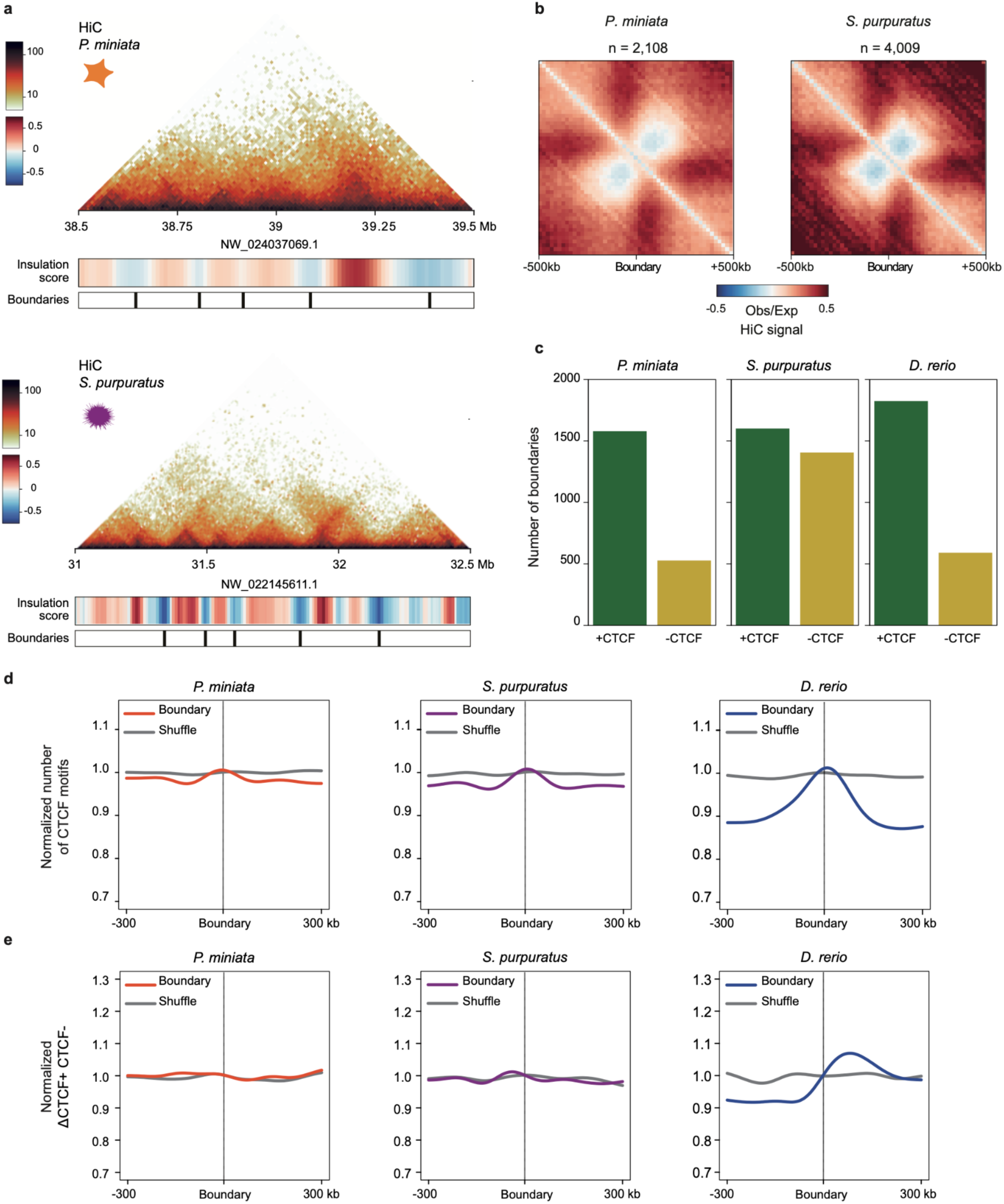
Echinoderm TAD boundaries and CTCF sites. **a**, Heatmaps showing normalized HiC signal at 10-kb resolution in *P. miniata* (top) and in *S. purpuratus* late gastrula embryos (bottom). Insulation scores and computationally called TAD boundaries are plotted below the heatmaps. **b**, Aggregate analysis of the observed versus expected HiC signal around TAD boundaries called at 10-kb resolution in *P. miniata* (left) and *S. purpuratus* late gastrula embryos (right). **c**, Number of boundaries harboring or not CTCF motifs. **d**, Average number of CTCF motifs within ±300 kb around boundaries (colored) and shuffled controls (grey). **e**, Normalized difference of CTCF motifs between forward and reverse strands within ±300 kb around boundaries (colored) and shuffled controls (gray).

### The Developmental Regulatory Genome of P. miniata and S. purpuratus

We next annotated the developmental regulatory genome of the two echinoderm species, identifying putative CREs (hereafter referred to as pCREs) as defined by profiling open chromatin regions with assay for transposase-accessible chromatin using sequencing (ATAC-seq). We performed ATAC-seq experiments in sea urchin and sea star whole-embryos at different timepoints ranging from blastula to larva, including three equivalent developmental stages between the two species (Fig. 1a) plus two additional stages in sea urchin, and identified a total of 66952 and 47919 pCREs in *P. miniata* and *S. purpuratus*, respectively (Fig. 3a, Supplementary Table 1). The genomic distribution of these elements was relatively similar between the two species, with the majority of pCREs (∼40%) laying in distal locations with respect to gene promoters or gene bodies (Fig. 3a). Sea urchin pCREs consistently overlapped with previously described CREs in this species, as in the case of the *S. purpuratus foxa1* gene, where we were able to identify pCREs that overlapped all four previously described regulatory elements in the intergenic regions surrounding it (Modules *F*, *I*, *J* and Region *K*) ^53^, as well as additional elements that had not been reported before (Fig. 3b). We tested seven of these *foxa1* pCREs in transient transgenic reporter assays ^54^ in sea urchin embryos. They drove GFP expression recapitulating the endogenous *foxa1* expression, both injected as pools, containing all seven pCRE-GFP constructs together, or individually (Fig. 3c). In the mesenchyme blastula, the construct pool drove GFP expression in the endoderm and oral ectoderm, with less than 3% of the injected embryos showing ectopic expression (mostly in the primary mesenchyme cells and misdeveloped embryos) (Extended Data Fig. 5a,c). Afterwards, in late gastrula, the expression was visible in the whole gut, equivalent to the expression patterns driven by previous construct concatenates (the FIJ CRE concatenate as described in ^53^ (Fig. 3c). Furthermore, quantitative expression assessment of the pCREs injected individually, showed that the sub-region K1 of the Region K, which was not previously analyzed in detail, was the strongest *foxa1* element tested, and its activity alone was enough to recapitulate all *foxa1* endodermal expression, consistent with the high accessibility of K1 in previously published ATAC-seq data from isolated gut samples (Extended Data Fig. 5b) ^49^. These results support that pCREs act as developmentally regulated transcriptional enhancers and are a good proxy for bona-fide CREs.

**Fig. 3:**
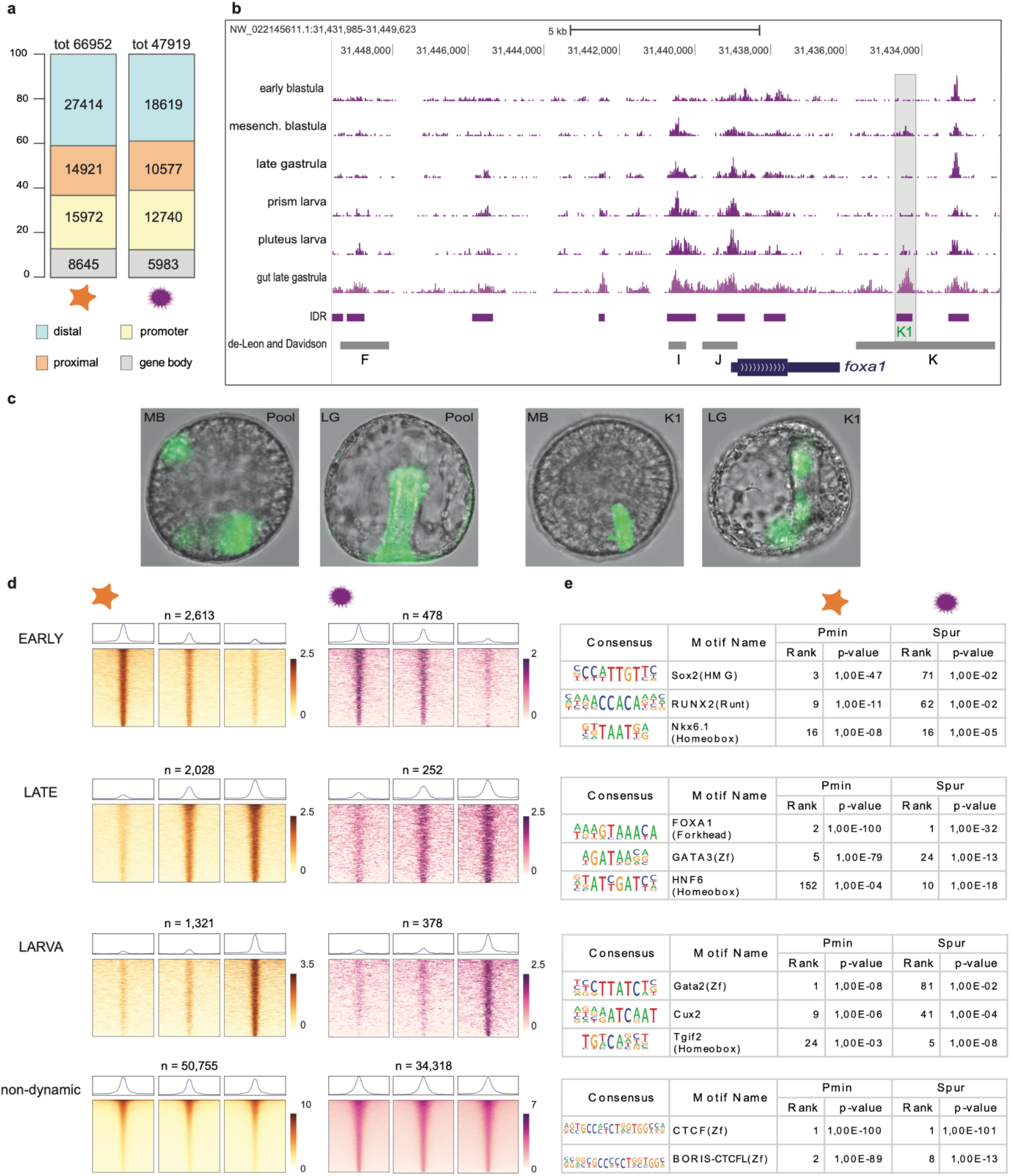
Regulatory annotation of echinoderm genomes using chromatin accessibility **a,** Numbers and proportions of open chromatin regions of both species according to their genomic location: promoters, for regions overlapping a gene TSS; proximal, within 5-kbp upstream of (but not overlapping with) a TSS; distal, not in the aforementioned categories. **b,** Screenshot of sea urchin UCSC genome browser with ATAC-seq tracks showing the correspondence between published ^53^ *foxa1* (LOC110977664) CREs (gray bars) and ATAC-seq peaks (pCREs, purple bars). **c,** From left to right, GFP expression driven by the *foxa1* pCREs pool (panels first and second) and the single K1 pCRE (third and fourth) at mesenchyme blastula (MB) and late gastrula (LG) stages. **d,** Heatmaps of clusters of pCREs of sea urchin (left panels, in purple) and sea star (right panels, in orange) that are more accessible at early (top panels), late (second row panels) and larva (pluteus/bipinnaria, third row panels) stages, as well as clusters of non-dynamic pCREs that did not show changes in accessibility across the sampled stages (bottom panels). Each heatmap column shows normalized ATAC-seq nucleosome-free signal for each stage ordered by developmental time (left: mesenchyme blastula, middle: late gastrula, right: bipinnaria/pluteus larva) over a 2 kb window centered in the midpoint of each of the pCREs, each heatmap horizontal line corresponds to a single pCRE. Average ATAC-seq signal profiles of all the pCREs in each stage are shown on top and the number of pCREs contained in each cluster are indicated at the lower right corner of each panel. **e**, Examples of TFBMs enriched in each of the temporal pCREs clusters that occupy top-ranked positions in both or just one of the two echinoderm species. The families of their corresponding TFs are indicated in brackets.

To better understand modules of regulation during development, we selected open chromatin regions that change their accessibility at least in one developmental stage, by clustering pCREs according to their signal profiles across the three matched developmental stages of the two species. Both in sea urchin and sea star, we identified three clusters of dynamic pCREs whose accessibility changes recapitulated the temporal progression across the three stages (“early”, “late” and “larva” clusters), as well as a fourth “non-dynamic” cluster which presumably contained constitutive pCREs (Fig. 3d). We then characterized the trans-regulation connected to these clusters of pCREs, by performing transcription factor binding motif (TFBM) enrichment analysis using a set of vertebrate TFBMs. These results showed clear similarities among the top-ranked TFBMs in the two species in some of the clusters (Fig. 3e, Supplementary Table 2). For instance, the motifs of Foxa genes were highly enriched in the late clusters of both species, highlighting the crucial role of this gene family in endoderm formation during gastrulation ^12,53,55–57^, while CTCF was the top-ranked motif of the non-dynamic clusters, consistent with a putative role of CTCF as an architectural protein, as suggested by its enrichment at TAD boundaries (Fig. 3e). However, differences in the temporal usage of TFBMs were much more prevalent, and the majority of top motifs in one species occupied much lower ranks in the other. This was the case of Sox motifs in early pCREs, which ranked much higher in sea star than in sea urchin, the motifs of several families of homeobox factors, such as Pitx and Otx, with more prominent positions in the early cluster of sea urchin, and Gata motifs, which were highly enriched in larva pCREs only in sea star (Fig. 3e, Supplementary Table 2).

### Regulatory conservation in echinoderms and deuterostomes

To study the evolution of sea urchin and sea star pCREs, we generated a multi-species alignment of their genomes together with additional genome sequences from multiple echinoderm and outgroup species, spanning a wide range of phylogenetic distances and divergence times (Fig. 4, Extended Data Fig. 6a,b, Supplementary Tables 3,4). This allowed us to detect blocks of conserved sequences between the genomes of the different species and, therefore, identify which sea urchin and sea star pCREs lay within these conserved blocks and determine their level of conservation. With this, we recognized four evolutionary strata (s) among the conserved regions. First, s1, the most recent ones, shared with other species of the same order/superfamily than our reference species (Valvatida for *P. miniata* and Odontophora for *S. purpuratus*) and dating back at least to ∼50-80 mya (^58^ Andrew Gale, personal communication). Second, s2, conserved with the most distantly related species within each of their respective echinoderm classes, Asteroidea and Echinoidea, that separated at least ∼200-265 mya ^59,60^. Third, s3, conserved across different echinoderm classes, including the two classes of our species as well as others such as Holothuroidea and Crinoidea, with divergence times of approximately 520 mya ^61^. And fourth, s4, conserved with other deuterostome phyla, such as hemichordates and chordates, which separated from echinoderms more than 550 mya ^62,63^. With this classification scheme, we assigned all sea star and sea urchin pCREs to a particular age class. For example, in the second intron of the sea star *hedgehog* gene (Fig. 4a), we found an Echinodermata pCRE, conserved across different echinoderm classes, plus another restricted to asteroidean species (i.e., an Asteroidea pCRE). Nearly 60% of sea urchin and sea star pCREs were not conserved with the other species, indicating a high degree of sequence turnover, as it has been previously reported in other animal lineages ^28,64,65^. Furthermore, among conserved pCREs (Fig. 4b and Extended Data Fig. 6b), the majority of them appear to have a relatively recent origin (∼50-80 mya), and are only conserved within Valvatida (23%) and Odontophora (36%), with a minimum age of a similar magnitude than the most ancient conserved CREs identified in nematodes and fruit flies, where only recent evolutionary elements are present ^30,32,33^. Remarkably, in stark contrast to these two ecdysozoan lineages, we also found that 16% (10588 pCREs) and 5% (2287 pCREs) of sea star and sea urchin pCREs were very ancient elements older than 200-265 my, predating the origin of the Echinoidea and Asteroidea crown groups in the late Permian and the Triassic (Fig. 4b). To date, this type of regulatory conservation, with several thousand elements that are at least several hundred my old, has only been reported in the vertebrate lineage ^39–43^. Finally, we also found a small number of pCREs with extremely deep conservation, shared by different classes of echinoderms (796 elements) or across other deuterostome phyla (208 elements) and whose origin dated back at least to the Cambrian, more than 500 mya ^66^. We next checked the genomic distribution of the different evolutionary strata of conserved pCREs. The most recent Valvatida and Odontophora elements were almost equally distributed across the different types of genomic locations, with a slightly higher prevalence in distal and proximal positions versus promoter locations (Fig. 4c, Extended Data Fig. 6b). The older class-specific Asteroidea and Echinoidea elements were also similarly prevalent at the different genomic locations, although in contrast to the previous age category, they were more frequently found at promoter positions. On the other hand, the most ancient pCREs, conserved between echinoderm classes and or deuterostome phyla, were mostly overlapping promoter regions. We then studied the degree of conservation of the temporal dynamic pCREs we identified in the previous section, as well as the non-dynamic, potentially constitutive, pCREs (Fig. 4d, Extended Data Fig. 6c). Interestingly, the conservation of developmental dynamic regions indicated that “early” and “larva” pCREs are the least conserved, showing the highest percentages of non-conserved species-specific elements, while the “late” pCREs were the most conserved in both species (Fig. 4d). “Non-dynamic” pCREs showed an intermediate pattern, with a lower percentage of conserved elements than the “late” clusters but higher than the “larva” and “early” clusters.

**Figure 4.**
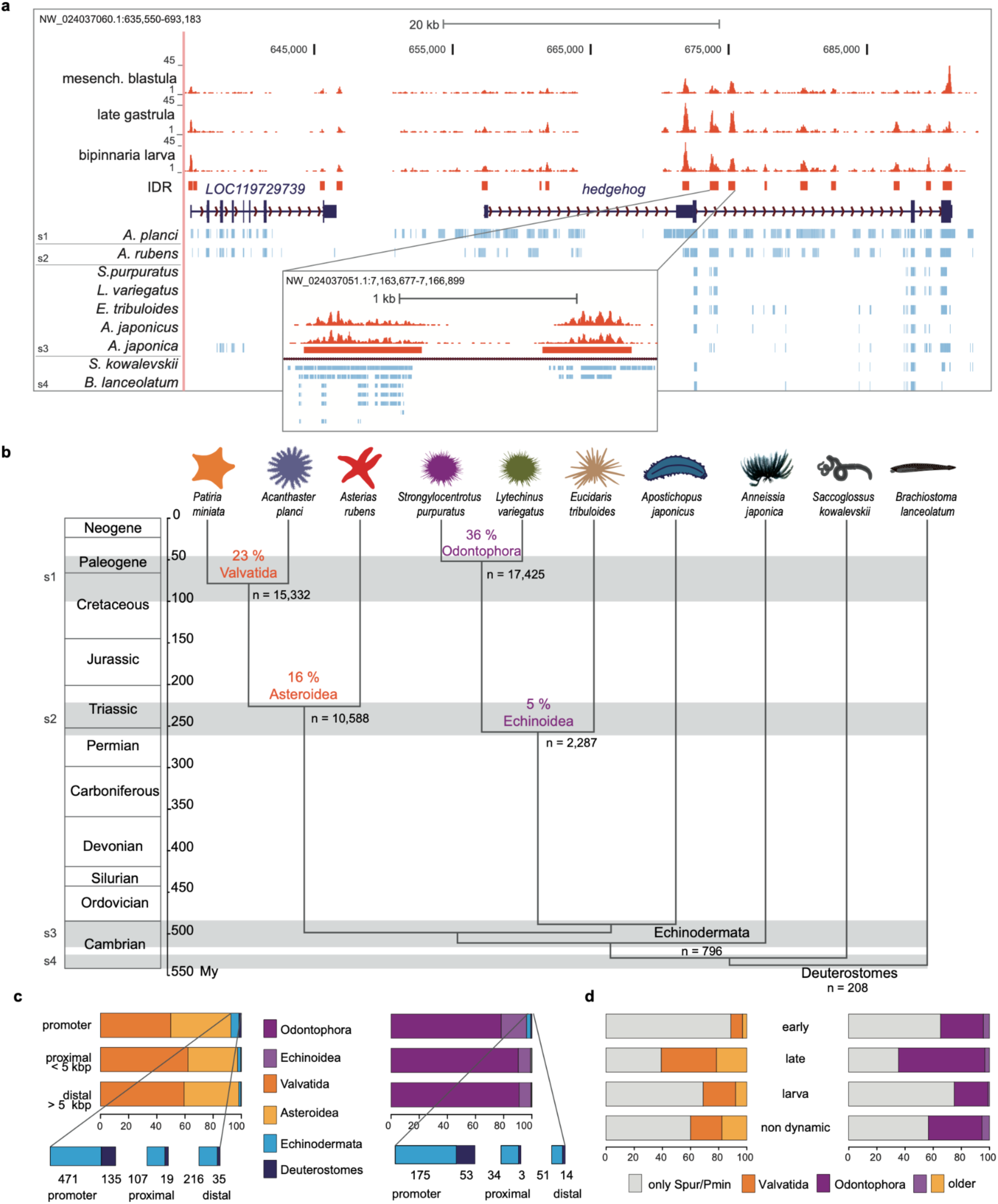
Evolutionary conservation of pCREs in echinoderms. **a**, Genome browser screenshot around the *P. miniata hh* gene (LOC119729696) showing ATAC-seq tracks from different developmental stages (orange) and the blocks of conserved sequences with other species included in the multiple genome alignment (light blue). The four evolutionary strata are indicated as “s1” to “s4”. Gene models are shown in dark blue. **b,** Schematic phylogenetic tree of the deuterostome species included in the genome alignments showing the percentage and total number of sea star (orange) and sea urchin (purple) conserved pCREs at each evolutionary strata (indicated by gray bars, s1 to s4). **c,** Genomic distribution of pCREs according to their evolutionary strata. Distributions are shown in orange for sea star and in purple for sea urchin. **d**, Proportions of conserved elements in each of the sea star and sea urchin pCREs temporal clusters (early, late, larva and non-dynamic).

Finally, we also checked whether the sea star and sea urchin deeply conserved transphyletic elements were also conserved regarding their chromatin accessibility by using previously published ATAC-seq data from one of the non-echinoderm phyla included in our analyses, an invertebrate chordate, the amphioxus *Branchiostoma lanceolatum* ^7^. We found a total of 78 sequence-conserved elements that were identified as accessible pCREs in both sea urchin and sea star ATAC-seq experiments, as well as a set of 16 elements that also have conserved chromatin accessibility with the chordate amphioxus (Fig. 5a, Supplementary table 5). We selected one of these 16 pCREs to perform functional assays in sea star and sea urchin embryos, a 5’ proximal element associated with Tbx2/3 genes, a family of transcription factors highly conserved in animals ^67^. This region had been identified as a deeply conserved element in previous works ^34,36^, and our results showed that it overlaps with ATAC-seq peaks in sea star, sea urchin and amphioxus (Fig. 5b,c). We cloned the two orthologous elements from sea star and sea urchin into GFP reporter vectors and assessed their gene regulatory activity by injecting them in embryos of the bat sea star, the purple sea urchin, and a second sea urchin species, *Paracentrotus lividus* (Fig. 5d, Extended Data Fig. 7a,b). The sea urchin Tbx2/3 endogenous spatiotemporal expression profile has been previously characterized and was found to be transiently expressed in several embryonic and larval domains. This includes both the skeletogenic and non-skeletogenic mesenchyme, dorsal regions of the gut, the anterior neuroectoderm, and the aboral/dorsal ectoderm, a domain in which it acts an early activator of the respective GRN ^68–70^. Similarly, in starfish embryos Tbx2/3 was previously found to be expressed in equivalent domains with the exception of the mesoderm ^71^. This is also corroborated by scRNA-seq data for both species (Extended Data Fig. 7c) ^72,73^. The GFP expression driven by the sea star and sea urchin Tbx2/3 pCREs in the two species of sea urchin embryos (Fig. 5d) matched the embryonic territories where the endogenous *Tbx2/3* gene is also expressed. Similarly, in the sea star embryos injected with either sea urchin or sea star constructs, GFP expression was also consistent with the endogenous *P. miniata Tbx2/3* expression described above (Fig. 5d). Thus, both the sea urchin and sea star Tbx2/3 CREs are able to drive the same expression patterns independent of the species of origin. Noteworthy, our results also indicate that Tbx2/3 expression in the aboral/dorsal ectoderm and the archenteron is conserved between these two distant echinoderm lineages (Fig. 5e). This showed that this ancient Tbx2/3 CRE is also conserved at the functional level.

**Figure 5.**
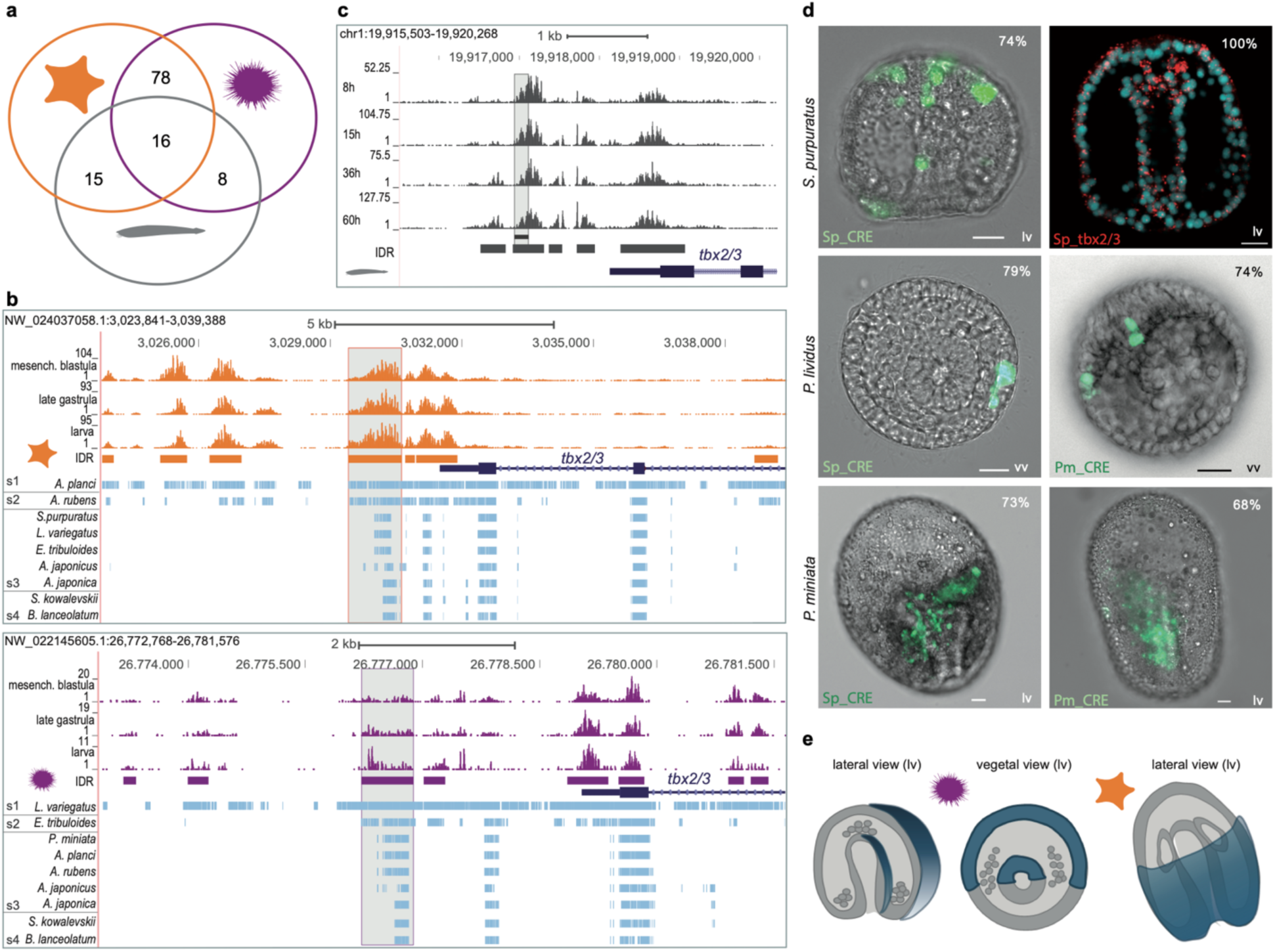
Deeply Conserved pCREs in deuterostomes. **a**, Venn diagram with the number of deeply conserved pCREs showing shared chromatin accessibility in sea star (orange), sea urchin (purple) and amphioxus (gray) embryos. **b,** Genome browser screenshots around the proximal promoter region of the Tbx2/3 genes of sea star and sea urchin. ATAC-seq tracks are colored in orange (sea star) and purple (sea urchin), gene models are shown in dark blue, and blocks of conserved sequences in the multiple genome alignment are indicated in light blue. In both screenshots the deeply conserved CRE tested in functional assays is highlighted by a light gray box. **c,** Genome browser screenshot around the proximal promoter region of the amphioxus *Tbx2/3* gene. ATAC-seq tracks are colored in gray and IDR ATAC-seq peaks are indicated by gray bars. The Tbx2/3 conserved sequence and the gene models are indicated as in (b). **d,** GFP expression driven by the *S. purpuratus tbx2/3* CRE in *S. purpuratus* (top left panel), *P. lividus* (central top panel) and *P. miniata* (right top panel) late gastrulae, and by the *P. miniata tbx2/3* CRE in *P. lividus* (central bottom panel) and *P. miniata* (right bottom panel) late gastrulae. Percentages represent the number of embryos showing expression in a specific domain compared to the total number of fluorescent embryos detected in the assay. Scale bar 20 µm. The GFP expression patterns are consistent with the endogenous expression of *tbx2/3* in the three species, as shown in the fluorescent *in situ* hybridization of *tbx2/3* in *S. purpuratus* late gastrula (top right panel) and as deduced from current (*S. purpuratus*) and previously published *in situ* hybridization data (*P. lividus*, ^74^) and analysis of scRNA-seq data (*P. miniata*, Extended Data Fig. 7 ^73^). **e,** Schematic drawings representing shared domains between sea urchin and sea star Tbx2/3 expression (in blue) in wild type and transgenic embryos.

## Discussion

By studying 3D chromatin organization and chromatin accessibility during echinoderm development, we have deepened our understanding of general trends in the evolution of the regulatory organization of animal genomes.

Our results added echinoderms to the list of animal lineages in which TAD-like structures are present ^21,75–82^. Our findings support the view that this is probably the ancestral condition in animals, as suggested also by multiple works linking the conservation of ancestral microsyntenic blocks with the presence of TADs ^3,14,15,17^, a link that we also observed in our work. From a more mechanistic perspective, the enrichments of CTCF motifs in echinoderm TAD boundary regions and in non-dynamic pCREs could be consistent with CTCF playing a role in chromatin architecture in the ancestor of bilaterian animals. However, the specific nature and relative importance of this putative ancestral role would be much more contentious. In contrast to vertebrates, where CTCF acts as a crucial factor in 3D chromatin and loop organization through its orientation-dependent binding ^18–21,83–85^ we did not find any obvious pattern in the orientation of CTCF motifs in sea star and sea urchin boundaries. Thus, in echinoderms the role of CTCF in chromatin folding could be more similar to that in fruit flies, where its function does not depend on orientation ^23^, suggesting that the vertebrate case is a derived condition that appeared later in evolution. Nevertheless, the derived features present in the CTCF proteins of echinoderms and other ambulacrarians, with the loss of ancestral Zn-finger domains and higher rates of evolution, would support an alternative scenario, with the independent loss of an ancestral vertebrate-like CTCF function in both flies and echinoderms. In any case, further work and data from additional animal lineages, will be needed to conclusively arbitrate between these different scenarios. In that regard, previous works in cephalopods and cephalochordates have reported CTCF motif enrichments in boundaries but without conclusively addressing the relative orientation of these motifs ^78,79^.

By focusing on echinoderms to study how regulatory elements evolve at multiple timescales, we have renewed long-standing debates about the biological significance of CRE conservation. We found that a large fraction of regulatory elements in echinoderms are evolving relatively rapidly. About half of sea star and sea urchin pCREs were not conserved at the sequence level in the rest of species we investigated. Furthermore, most of the rest of the elements were young and only conserved with close relative species. This is consistent with previous works addressing non-coding sequence conservation between *S. purpuratus* and *L. variegatus* sea urchins as well as in other lineages ^26,30–33,53,86–90^. These observations suggest that a high rate of CRE turnover is a shared feature across all studied species, and a universal property of the evolution of gene regulation in animals. In contrast, a high prevalence of regulatory sequences of very ancient origin is a much more puzzling phenomenon. Until now, the several thousand elements, older than 440 my, that are shared across jawed vertebrates ^39,40,42^ had seemed to be an exclusive feature of the vertebrate lineage. Our results have completely changed this picture. We have found thousands of deeply conserved pCREs (from 2,287 in sea urchin to 10,588 in sea star) that date back at least to the late Triassic or even the Permian (200-265 mya), showing that this conservation pattern, previously considered unique to vertebrates, is also present in echinoderms. In fact, it would be more appropriate to say that both vertebrates and echinoderms share an ancestral feature of regulatory evolution that was already present in deuterostome ancestors.

A small fraction of elements, a few hundred pCREs, was conserved even deeper, between echinoderm classes or even across deuterostome phyla. The fact that we found a similarly low amount of transclass and transphyletic conserved pCREs in echinoderms suggests that, from the perspective of cis-regulatory conservation, the different classes of echinoderms were almost as divergent as different deuterostome phyla. This divergence was also reflected in terms of TFBM usage, with pronounced differences in TFBM enrichments between sea star and sea urchin pCREs associated with equivalent developmental stages. Nevertheless, the extreme conservation of the few transclass and transphyletic pCREs we identified, especially the set of elements with conserved chromatin accessibility, such as the Tbx2/3 enhancer we functionally tested, suggest that they may be particularly important for developmental and evolutionary aspects of gene regulation shared by the echinoderm and chordate phyla. In this regard, the higher conservation of pCREs that increase their accessibility during the late gastrula stage would be consistent with previous studies comparing different species of sea urchins and of additional animal lineages, where mid-developmental stages show the highest degree of morphological and gene expression conservation ^7,12,91–95^.

Low numbers of extremely old “Cambrian-age” transphyletic conserved non-coding elements have also been reported in some species outside of deuterostomes, including certain species of molluscs, brachiopods, chelicerates, priapulids and cnidarians ^34–36^. Typically, many of these species also share with echinoderms other hallmarks of relatively slow genome evolution, such as a high level of conservation of ancestral microsyntenic blocks around key developmental genes and high retention of ancestral gene families ^13,16,45,46^. Thus, although these transphyletic elements probably constitute just a very tiny fraction of the total amount of CREs present in any of these species, our results in echinoderms suggest that they can actually be the smoking gun indicating that a vertebrate-like pattern of regulatory conservation is also present in these additional animal lineages. Indeed, recent results studying non-coding conservation in mollusks, cnidarians and arthropods point into this direction, identifying a high number of ancient conserved non-coding sequences in some of these lineages ^96^. If these conserved sequences were confirmed as CREs by functional genomics data, it would indicate that the conservation of thousands of ancient CREs, as in the vertebrate and echinoderm cases, could be an ancestral property of the regulatory organization of animals, paving the way to finally understand the so far elusive evolutionary and functional causes underlying CRE sequence conservation.

## Methods

### Animal origin, gamete collection, and in vitro fertilization

Sea urchin *Strongylocentrotus purpuratus* and sea star *Patiria miniata* animals were provided by Patrick Leahy (Kerckhoff Marine Laboratory, California Institute of Technology, Pasadena, CA, USA) and kept in a closed tank system with circulating diluted Mediterranean Sea water at 36 ppt and 14 °C at Stazione Zoologica Anton Dohrn. Sea urchin gametes were obtained by vigorously shaking the adults. Eggs were collected by placing the spawning female over a beaker with filtered sea water (fSW) placed on ice, with the aboral side up so that the animal would be partially submerged. Sperm was collected by pipetting from the surface of the spawning male. Gametes of sea star were obtained by making a V-shaped surgical incision on the aboral side of the animal next to the gonads. The female and gonads were torn up under the dissecting microscope to release gametes into the fSW. Eggs of both species were then passed through a 200 μm Nitex mesh. Sea star oocytes were also treated with 10 µM 1-methyladenine after passing through a filter to mature, until the germinal vesicle disappears, after which eggs were washed with fSW. Diluted sperm (5 μl of dry sperm in 13 ml of fSW) was added drop-wise to fertilize the eggs. Fertilized eggs were cultured in fSW at 15°C until the desired stage.

### Genome Sequencing and Contig Draft Assembly

*P. miniata* genomic DNA was obtained anew from the gametes of a single male animal, while *S. purpuratus* was sequenced using the preserved genomic DNA samples from the same single male preparation used by the Sea Urchin Genome Sequencing Consortium for the first draft of the purple sea urchin genome assembly ^45^. High molecular weight DNA larger than 50 kbp was selected using the Circulomics Nanobind CBB Big DNA Kit. The isolated DNA was approximately 300 ng/ul and the majority was between 48 kbp and 145 kbp, as verified by a clamped homogeneous electric field (CHEF) gel.

Approximately 30 ug of the high molecular weight genomic DNA from each species was sent to the Duke University School of Medicine Sequencing and Genomic Technologies facilities, where they performed large insert library prep and PacBio Sequel sequencing with 16 Single Molecule Real Time (SMRT) cells.

The PacBio Sequel reads were assembled into contigs using Canu v1.8 for *S. purpuratus*, and Canu v. DEC-2019 for *P. miniata* ^97^ with default parameters. To assess the estimated completeness of the draft assembly, we performed a Benchmarking Universal Single-Copy Ortholog (BUSCO) ^98^ analysis for both species using a set of 978 metazoan genes.

### HiC Sequencing and Alignment of HiC Reads

*P. miniata* embryos were fixed in 2% paraformaldehyde in phosphate-buffered saline (PFA/PBS) at 68 hpf. *S. purpuratus* embryos were fixed in 2% PFA/PBS at 50 hpf. The fixation reaction was quenched after 10 minutes of incubation by adding glycine to a final concentration of 0.125 M. Subsequently embryos were washed twice with 1x PBS. The fixed embryos were frozen in liquid nitrogen and sent to the Gómez-Skarmeta lab for HiC library preparation and sequencing. HiC was performed in duplicates for *P. miniata* and *S. purpuratus*, with about 10 million cells per replicate. HiC preparation was based on ^18^ with minor modifications, described in detail in ^21^. Briefly, nuclei were isolated and permeabilized by SDS treatment. Chromatin was fragmented using the DpnII restriction enzyme (NEB, R0543). Restriction overhangs were blunted and marked with biotin-14-dATP and subsequent proximity ligation was performed in intact nuclei. Purified and sonicated biotin-labeled DNA fragments were enriched using Dynabeads My One C1 Streptavidin beads. Sequencing library preparation and sample indexing were based on TruSeq Illumina technology. Final library for paired-end sequencing was prepared using NEBNext High-Fidelity 2X PCR Master Mix (NEB), using TruSeq Primer 1.0 (P5) and TruSeq Primer 2.0 (P7). For each sample and replicate, at least eight independent PCR reactions were performed and then pooled to maintain library complexity for sequencing. Libraries were multiplexed and sequenced using DNBseq technology (BGI, China) to produce 50 bp paired-end reads and approximately 400 million raw sequencing read pairs per sample.

HiC paired-end reads were mapped to the *P. miniata* and *S. purpuratus* genome assemblies using BWA ^99^. Reads from biological replicates were pooled before mapping. Then, ligation events (HiC pairs) were detected and sorted, and PCR duplicates were removed, using the pairtools package (https://github.com/mirnylab/pairtools). Unligated and self-ligated events (dangling and extra-dangling ends, respectively) were filtered out by removing contacts mapping to the same or adjacent restriction fragments. The resulting filtered pairs file was converted to a tsv file that was used as input for Juicer Tools 1.13.02 Pre ^100^, which generated multiresolution hic files. These analyses were performed using custom scripts (https://gitlab.com/rdacemel/hic_ctcf-null) ^23^: the hic_pipe.py script was first used to generate tsv files with the filtered pairs, and the filt2hic.sh script was then used to generate Juicer hic files. HiC matrices at 10 Kb resolution, normalized with the Knight-Ruiz (KR) method ^101^, were extracted for downstream analysis using the FAN-C 0.9.14 toolkit ^102^. Visualization of normalized HiC matrices and other values described below, such as insulation scores, TAD boundaries, aggregate TAD and loop analysis, were calculated and visualized using FAN-C.

TAD boundaries were called using the insulation score method ^103^. Insulation scores were calculated for 10-Kb binned HiC matrices using FAN-C ^102^. Briefly, the average number of interactions of each bin were calculated in 250-Kb square sliding windows (25 x 25 bins); then, these values were normalized as the log2 ratio of each bin’s value and the mean of all bins to obtain the insulation score for each bin; next, minima along the insulation score vector were calculated using a delta vector of +/-50 Kb (+/-5 bins) around the central bin. Only boundaries with a score lower than 1.5 were considered, to avoid calling low mappable regions as TAD boundaries. Chromatin loops were called using Juicer Tools 1.13.02 Hiccups ^18^, with standard parameters. Briefly, the multiresolution hic file was used as input for the CPU version of HICCUPS, which ran using 5, 10 and 25-Kb resolution KR-normalized matrices. The maximum permitted FDR value was 0.1 for the three resolutions; the peak widths were 4, 2 and 1 bin for 5, 10 and 25-Kb resolutions, respectively; and the window widths to define the local neighborhoods used as background were 7, 5 and 3 bins, respectively. The thresholds for merging loop lists from different resolutions were the following: maximum sum of FDR values of 0.02 for the horizontal, vertical, donut and lower-left neighborhoods; minimum enrichment of 1.5 for the horizontal and vertical neighborhoods; minimum enrichment of 1.75 for the donut and bottom-left neighborhoods; minimum enrichment of 2 for either the donut or the bottom-left neighborhoods. The distances used to merge the nearby pixels to a centroid were 20, 20 and 50-Kb for 5, 10 and 25-Kb resolutions, respectively. Only loops whose anchors were located in the same contig were considered.

### Creating Scaffolds

The draft contigs for each species were assembled into scaffolds using the 3D de novo assembly (3D-DNA) v.180922 pipeline (https://github.com/aidenlab/3d-dna) ^104^. This pipeline uses the HiC alignment data generated by Juicer to construct scaffolds based on 3D proximity relationships. These scaffolds and HiC contacts can be viewed as heatmaps using the Juicebox visualization tool (https://github.com/aidenlab/Juicebox) ^105^.

The draft contigs were assembled into scaffolds with the 3D-DNA *run-liger-scaffolder* program, using contigs larger than 6 kbp and HiC reads with a MAPQ alignment score greater than 0. Inspecting the HiC contact heatmap revealed a large number of overlapping contigs, which appear as parallel or perpendicular lines of HiC contacts between two contigs. These overlaps were dealt with using the 3D-DNA *align-nearby-sequences-and-filter-overlaps* and *tile-assembly-based-on-overlaps* programs. This identified and merged overlapping contigs within the scaffolds, reducing the size of the draft assembly from 1075 Mbp to 714Mbp. The finalized assemblies were submitted to the NCBI genome database GenBank, where they are publicly available with accession numbers GCA_015706575.1 (*P. miniata* v3.0) and GCA_000002235.4 (*S. purpuratus*).

### Annotation of the assemblies

Assemblies were annotated using the NCBI eukaryotic genome annotation pipeline ^51^ using publicly available Entrez transcripts and RNA-seq reads from the Sequence Read Archive (SRA). Gene content was also evaluated by using BLAST ^106^ to align the previous assembly’s gene models to the new assembly.

### Generation of cohesin (SA1-3) and CTCF gene trees

Protein sequences of cohesin subunit SA1-3 and of CTCF were searched using the Blast server (https://blast.ncbi.nlm.nih.gov/Blast.cgi), restricting by taxonomy to specifically search in the genomes and proteomes of those animal lineages of particular relevance for the phylogenetic context of echinoderms. *Mus musculus* and *Drosophila melanogaster* protein sequences were used as initial queries. After an initial search, a second one was performed using the retrieved protein hits as new queries. In the case of *Hemicentrotus pulcherrimus* SA1-3 protein, the sequence was obtained from transcriptome shotgun assembly data.

For the phylogenetic analyses, protein sequences were aligned with the MAFFT software ^107^ and the resulting alignments were trimmed (Supplementary Data 1,2) using Aliview to discard spuriously aligned regions ^108^. Phylogenetic trees were built using IQ-Tree ^109^, testing the tree with UFBoot (bootstrap = 10^3^) ^110^ and an approximate Bayes test for single branch testing. Following previous CTCF phylogenetic studies ^111^, the tree was rooted using the CROL proteins of *Drosophila melanogaster* and *Tribolium castaneum*. In the case of SA1-3 proteins, the tree was rooted using orthologs from sponges. The model used (LG+G4+I for both trees) was selected using ModelFinder with BIC as the criteria^112^. The trees (Supplementary Data 3,4) were visualized using ITOL ^113^.

### Multiple Genome Alignments

A multiple genome alignment was constructed using cactus v1.2.3 ^114^ with default parameters. This contained our two newly reported genomes, a broader sampling of echinoderms, and two marine deuterostomes (*Branchiostoma lanceolatum* and *Saccoglossus kowalevskii*). A full list of included species and genome accession numbers can be found in Extended Data Fig 6a. The resulting .hal files are hosted on the Echinobase ^115^ FTP server (https://download.xenbase.org/echinobase/Genomics/user-submitted/MGA_echinoderms/NewMGA/FinalMGA/), and associated .hub files to describe the alignment properties were generated and formatted per the UCSC Genome Browser User Guide. The alignments can be viewed via the UCSC Genome Browser sessions https://genome.ucsc.edu/s/echinoreg/Pmin for alignments anchored on *P. miniata* genome and https://genome.ucsc.edu/s/echinoreg/Spur for ones anchored on *S. purpuratus* v5.0 genome assembly (permanent links TBD).

### ATAC-seq library preparation

ATAC-seq libraries were generated as described in ^116,117^. Cultured embryos were manually collected into 1.5 ml Eppendorf tubes with the aim to get around 50,000 cells. The embryos were then centrifuged at 500 x g to remove liquid, and washed with fASW twice. The embryos were then lysed in 50 μl of lysis buffer (10 mM Tris pH 7.4, 10 mM NaCl, 3 mM MgCl2 and 0.2% NP40/Igepal CA-630). The half of nuclei suspension were incubated for 30 minutes at 37°C with 25 μl of 2x tagmented DNA buffer (TD buffer: 20 mM Tris(hydroxymethyl)aminomethane; 10 mM MgCl2; 20% (vol/vol) dimethylformamide) and 1.25 μl Tn5 enzyme and Tagmentation Buffer (10 mM Tris–HCl pH 8.0, 5 mM MgCl2, 10% w/v dimethylformamide), and incubated for 30 min at 37°C. Then, tagmented DNA was purified with the Qiagen MinElute kit (Qiagen, 28004) and eluted in 10 μl. PCR reactions for each replicate were performed with 15 cycles using Ad1F and different AdR primers ^116^) and NEBNext High-Fidelity 2X PCR Master Mix (New England Labs Cat #M0541). The resulting libraries were multiplexed and sequenced on the HiSeq 4000 sequencer.

### ATAC-seq data analysis

Data analysis for ATAC-seq libraries were performed as described in ^117^. After sequencing, paired-end reads without adapter sequences were aligned against the reference genome using Bowtie2 v2.2.6 ^118^ with *-X 2000-no-mixed-no-unal* parameters, allowing the retention of reads that are separated < 2 kb. PCR artifacts and duplicates were removed using the tool *rmdup*, available in the Samtools v1.3 toolkit ^119^. Read start sites were offset by +4 or by −5 bp in the plus or minus strand, respectively. Nucleosome-free pairs (insert < 130 bp) retained for peak-calling using MACS2 v2.1.1.20160309 ^120^ with the following parameters: *–nomodel –shift −45 –extsize 100* and the genome size of the correspondent organism. The irreducible discovery rate (IDR v2.0.3, ^121^) was used to assess replicability with the following parameters: *–input-file-type narrowpeak –rank p.value –soft-idr-threshold 0.1* and the MACS2 peak calling. Called IDR peaks were split into promoter, gene body and noncoding regions following genomic position by using the aforementioned gene annotation as reference. Promoters were defined as those ATAC peaks overlapping a single annotated TSS of genes; proximal noncoding locations within 5-kbp upstream of a TSS without overlapping with it and distal noncoding locations, not in the aforementioned categories and gene body. For conservation analyses, in ATAC peaks overlapping exon1 in promoter regions, only noncoding sequences were considered. The proportion of reads inside ATAC peaks were calculated with intersectBed from BEDtools v2.26.0 ^122^, employing the parameter *-c* where the reference was the high confidence set of ATAC peaks and the query was the filtered reads. The resulting counts table was then used in the DESeq2 v1.18.0 R package ^123^ and those ATAC peaks with a *p*-value under 0.05 were selected as differentially accessible ATAC peaks among developmental stages. All ATAC peaks as well differentially accessible peaks were associated with their putative target genes using the closestBED tool utility from BEDtools, with the parameters *-D ref -iu -no-namecheck -k 1*. The output file was used to calculate the percentage of ATAC peak distribution. In order to find TFBM enrichments in ATAC peaks was used the script FindMotifsGenome.pl from HOMER v3.12 software ^124^, selecting the set of desired TFBMs with the parameter *–mset* and using, merge high confidence ATAC peaks as background model. To find specific temporal dynamics in open chromatin regions, dynamic ATAC peaks were analyzed with seqMINER v1.3.4 ^125^ and cluster heat map plots were generated by computeMatrix deepTool v3.4.3 ^126^. The .hal files from multiple alignment analysis were processed with ATAC-seq IDR files of different genomic positions to return a correspondence between ATAC peaks of query species and a reciprocal mapping has at least 50% overlapping with the original ATAC peak coordinate, taken out with halLiftover v2.1 tool ^127^ with the option *--noDupes*. IntersectBed form BEDtool, with the options -f 0.5 -r or -f 0.5 -r -a, were used to cross genomic position between species. Resulting files were intersected with called ATAC peaks and those lists were grouped following their relative evolutionary distance from the query species by using *subset()* and *merge()* R basic command (https://www.R-project.org/).

### CRE DNA tag synthesis

Tagging of putative CREs was performed according to the protocol described by ^54^. Primers for the putative CREs were designed using Primer3web 4.1.0 ^128^ additional 18bp of the reverse complement of the beginning of the DNA tag sequence added to the 5’ of the reverse primer to ensure that the CRE can be combined with the DNA ^129^. Equal amounts of CRE and Tag DNA were combined using overlap PCR ^54,129^ using Expand High Fidelity PLUS PCR (Sigma). The product was gel purified using 2% agarose 1X TAE gel and GenElute Gel Extraction Kit (Sigma) (FoxA pCREs) or NucleoSpin Gel & PCR Clean-up (Macherey-Nagel) (Tbx2/3 pCREs) according to manufacturer’s guidelines and eluted in 50 μl of Elution buffer. The eluted DNA was then purified again using QIAquick PCR Purification Kit (Qiagen) and eluted in 30 μl of elution buffer. NanoDrop ND-1000 (ThermoFisher Scientific) was used to assess DNA yield and quality.

### CRE Microinjections

Gametes for sea urchin and sea star were collected as described above. Sea urchin eggs were de-jellied and placed on 4% protamine sulfate (PS) treated plates and then fertilized, while the sea star ones were first fertilized, de-jellied and then placed on the PS treated plates. Sea urchin and sea star zygotes were injected with 2 pL and 8 pL of microinjection solution, respectively. Microinjection solutions consisted of 0.5 μl of tagged CRE pool, 1.2 μl of 1mM KCl, 0.275 μl of 500 ng/μl carrier DNA and water up to 10 μl ^54,130^. The product of CRE tagging was used for microinjections either individually or in a pool. Injected eggs were then washed twice with fSW and incubated at 15℃ overnight for *S. purpuratus* and *P. miniata* and at 18℃ for *P. lividus*. Upon hatching embryos were transferred to 4-well plates (ThermoFisher Scientific) with fSW to grow until the desired stage.

### Genomic DNA and mRNA extraction from CRE microinjected embryos

Microinjected embryos of the desired stage were collected into 1.5 ml tubes (Eppendorf). AllPrep DNA/RNA Micro kit (Qiagen) kit was used to extract genomic DNA and RNA from them according to manufacturer’s guidelines. DNA was eluted in 100 μl of 65°C nuclease free water, RNA was eluted in 18 μl of 65°C nuclease free water. The RNA was treated with DNase from RNAqueous-Micro Total RNA Isolation Kit (Invitrogen) according to manufacturer’s guidelines. 14 μl of the DNase treated RNA was used to synthesize cDNA using SuperScript VILO cDNA Synthesis Kit (Invitrogen) as per manufacturer’s guidelines. The synthesized cDNA and eluted DNA was used for qPCR quantification.

### qPCR quantification of CRE expression

Extracted genomic DNA and synthesized cDNA were used to estimate relative expression of the microinjected tagged CREs in two biological replicates. Prior to qPCR the cDNA was amplified using universal primers ^54^ using the following thermocycler program: 2 minutes at 95°C, 21 cycles of 15 seconds at 95°C, 30 seconds at 60°C and 1 minute at 72°C, followed by 5 minutes at 72°C and hold at 4°C. The product was purified using QIAquick PCR Purification Kit (Qiagen) and eluted in 30 μl and used for qPCR quantification: 5 μl of Fast SYBR Green Master Mix, 4 μl of 0.7 pmol/ul qPCR primers and 1 μl of cDNA/gDNA. ViiA7 Real–Time PCR System machine (Life Technologies) was used for 20 seconds at 95°C, 40 cycles of 1 second at 95°C and 20 seconds at 60°C, followed by melting curve at 95°C for 15 seconds, 60 °C for 1 minute and 95°C for 15 seconds to perform the quantifications. Results were collected and exported into csv files. Total GFP expression was used as control for quantification of cDNA and genomic samples with the same specific tag primers. The number of tags expressed was normalized to the number of tags in the genomic DNA by dividing the number of expressed tags in cDNA by the number of expressed tags in gDNA relative to GFP. Mean of the between-replicate values was used for plotting.

### CRE expression visualization

Microinjected embryos of the desired stage were collected onto a two-well slide. A few drops of 100% methanol were added to immobilize the embryos. Embryo GFP fluorescence was visualized using Zeiss Imager.Z2 using GFP and bright-field settings^130^. Images with split channels were merged into a single image per embryo using ImageJ 1.52o.

### Fluorescent in situ hybridization (FISH)

FISH for Sp-Tbx2/3 gene was performed as previously described ^131^. The primes used to isolate the Sp-Tbx2/3 cDNA from gastrula stage total cDNA are described in Valencia et al., 2021. Digoxigenin-labelled antisense RNA probe for Sp-Tbx2/3 was synthesized as previously shown ^132^. Briefly, *S. purpuratus* gastrula stage embryos (48 hpf) were collected and fixed for 1h at RT in 4% PFA in MOPS buffer (0.1 M MOPS pH 7, 0.5 M NaCl and 0.1% Tween-20 in nuclease-free water). Embryos were washed several times with MOPS buffer and stored in 70% ethanol at −20℃. After rehydration, embryos were incubated with the probe solution in Hybridization buffer (50% formamide, 0.1 M MOPS pH 7, 0.5 M NaCl and 0.1% Tween-20, 1 mg/ml Bovine serum albumin (BSA) in nuclease-free water) at 65℃ overnight. The following day the probe solution was removed, the embryos were washed several times with MOPS buffer, incubated with blocking solution (Akoya Biosciences) for 30 min and with anti-digoxigenin POD (Roche) for 1 h and 30 min at 37℃. The fluorescence signal was developed using the fluorophore-conjugated tyramide technology using TSA Plus Cyanine 3 and 5 kits (Akoya Biosciences). Embryos were mounted and imaged using a Zeiss LSM 700 confocal microscope.

## Data availability

Genome assemblies used in this work are publicly available at the NCBI genome database GenBank with accession numbers <u>GCA_015706575.1</u> (*P. miniata*) and <u>GCF_000002235.5</u> (*S. purpuratus*). The HiC (<u>GSE281901</u>) and ATAC-seq (<u>GSE280529</u>) raw and processed sequencing data were deposited at the Gene Expression Omnibus (GEO) database under accession code of the SuperSeries <u>GSE281904</u>, with the exception of ATAC-seq data in *S. purpuratus* at 48 hpf that were deposited in <u>GSE186363</u>.

## Acknowledgements

JLG-S was supported by the European Research Council (grant agreement no. 740041) and the Spanish Ministerio de Economía y Competitividad (grant no. PID2019-103921GB-I00). VH, ST, GAC, and CK were supported by grant P41HD09583106 funded by the NIH. DV, MLR and CC were supported by the Stazione Zoologica Anton Dohrn PhD fellowships. This work was supported by the H2020 Marie Skłodowska-Curie Actions Innovative Training Network EvoCELL (grant number 766053 to MIA and fellowship to PP). MSM was granted a fellowship of the Program for the Training of Researchers (BES-2014-068494) and MFM holds a FPU fellowship (FPU20/02733), funded by the Spanish Ministry of Science, Innovation and Universities (MICIU). IM, MSM and MFM are supported by grants PID2021-128728NB-I00 and CNS2022-136105 funded by MICIU/AEI/10.13039/501100011033 and by “ERDF/EU” and “European Union NextGenerationEU/PRTR”. IM was also funded by grants RYC-2016-20089 and PGC2018-099392-A-I00 funded by MICIU/AEI/10.13039/501100011033, “ERDF A way of making Europe” and “ESF Investing in your future”. The authors thank Demian Burguera and Isabel Almudi for helpful discussions.

## Author Contributions

DNA library preparation: GAG; *S. purpuratus* genome assembly: GAG, SF; *P. miniata* genome assembly: CK, SF; Gene annotation coordination: SF; ATAC-seq experiments: MSM, DV, JR, CC; ATAC-seq data analyses: MSM; TF motif analyses: MSM, PF, PMM-G; HiC experiments: MF, RDA; preparation of samples for HiC: DV, CC, PP; HiC data analyses: JMS-P, PMM-G; gene family evolution analyses: MFM; multi-species genome alignments: SF; AG-G; sequence conservation analyses: MSM, AG-G; preparation of constructs and *cis*-regulatory reporter assays: DV, PP, MLR, MF-M; microinjection experiments: MIA; whole mount in situ hybridization: PP; conceptualization and design of the project: IM, MIA, JLG-S, VH; project coordination and supervision: IM, MIA, VH; funding acquisition: JLG-S, VH, MIA, IM; figure preparation: MSM, DV, SF, PMM-G, JMS-P, MF-M, PP, MLR, IM, MIA; manuscript writing – original draft: MSM, DV, SF, IM; manuscript writing – review & editing: IM, DV, MSM, SF, MIA, VH, CK, JMS-P, PF, RDA, PP, MF-M. All authors revised and approved the manuscript.

## Ethics declarations

Competing interests:

The authors declare no competing interests.

## Image Credits

Animal silhouettes from *A. planci*, *P. miniata* and *A. japonica* were drawn by I.M. under a CC0 1.0 Universal Public Domain Dedication. *B. lanceolatum* drawing is a black-colored modified version of an illustration by Juan J. Tena under a Creative Commons Attribution (CC-BY) license. The rest of silhouettes were obtained from Phylopic (https://www.phylopic.org/), with the following credits: *A. rubens* was represented by a red-colored modified version of the Asteriidae silhouette by Mali’o Kodis, photograph by “Wildcat Dunny” (http://www.flickr.com/people/wildcat_dunny/) under an Attribution 3.0 Unported license (https://creativecommons.org/licenses/by/3.0/); *S. kowalevskii* was represented by a *Balanoglossus* silhouette by Yan Wong; *Ap. japonicus* was represented by a blue-colored modified version of *Holothuria* silhouette by Lauren Sumner-Rooney; *E. tribuloides* was represented by a beige-colored modified version of *E. tribuloides* silhouette by Jake Warner; and *S. purpuratus* and *L. variegatus* were represented by purple and green-colored modified versions of *S. purpuratus* silhouette by Christoph Schomburg, all four of which (*Balanoglossus*, *E. tribuloides*, *Holothuria*, and *S. purpuratus*) released under CC0 1.0 Universal Public Domain Dedications (https://creativecommons.org/publicdomain/zero/1.0/).

## Supplementary Information

**Supplementary Data 1:** MAFFT alignment of SA1-3 proteins

**Supplementary Data 2:** MAFFT alignment of CTCF and CROL proteins

**Supplementary Data 3:** SA1-3 tree

**Supplementary Data 4:** CTCF tree

**Supplementary Table 1:** ATAC-seq statistics

**Supplementary Table 2:** Selected TFBM enrichments in “early”, “late” & “larva” pCREs

**Supplementary Table 3:** Conservation of P. miniata pCREs

**Supplementary Table 4:** Conservation of S. purpuratus pCREs

**Supplementary Table 5:** Deeply conserved pCREs in sea star, sea urchin and amphioxus

## Extended Data Figures

**Extended Data Figure 1.**
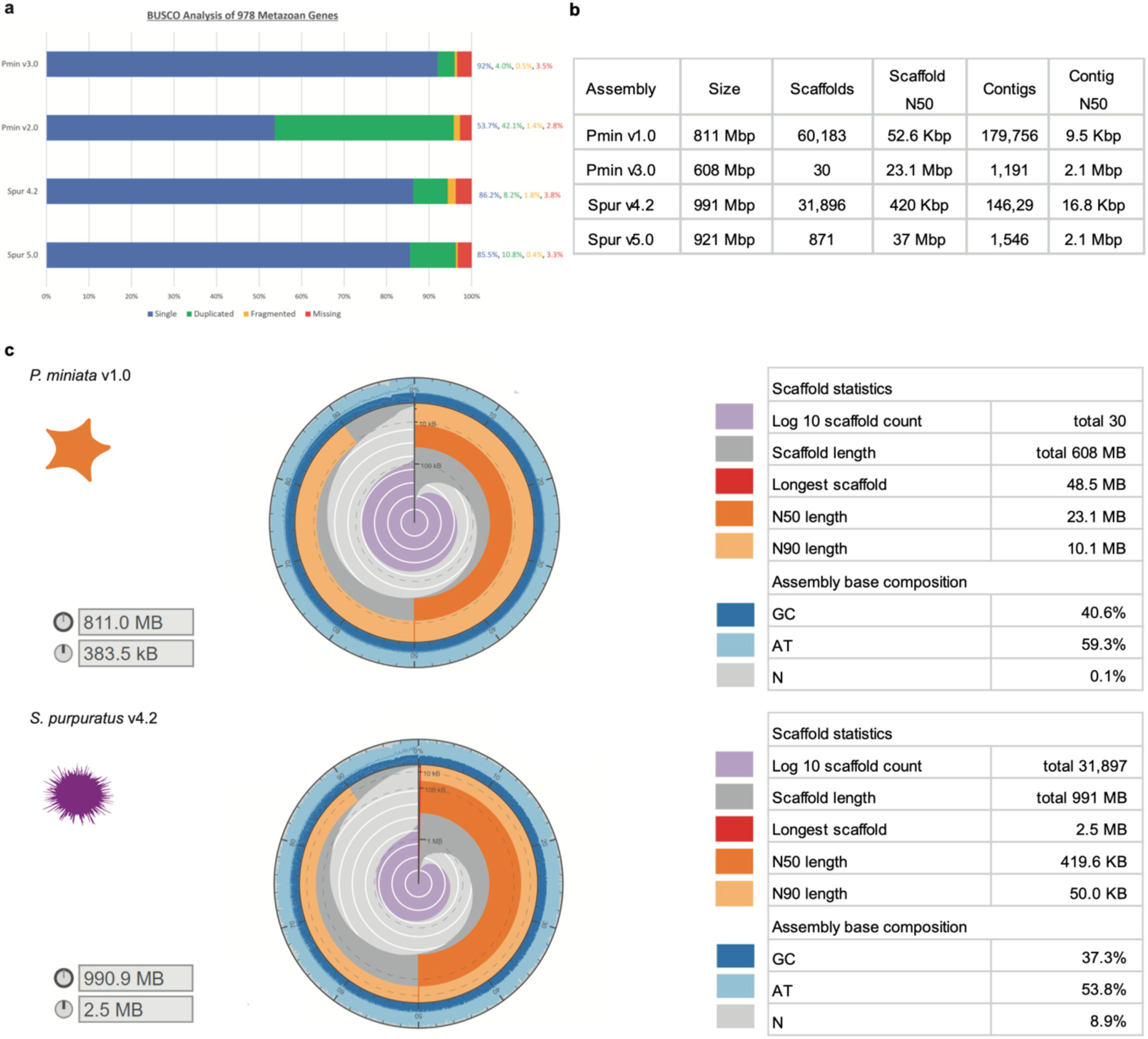
Comparison of current and previous assembly statistics: **a**, BUSCO statistics of current and previous genome assemblies. **b**, Assembly statistics of the two new genomes along with their older versions for comparison, generated with the assembly-stats perl package. **c**, Snailplots of general metrics of previous assemblies of purple sea urchin and bat sea star.

**Extended Data Figure 2.**
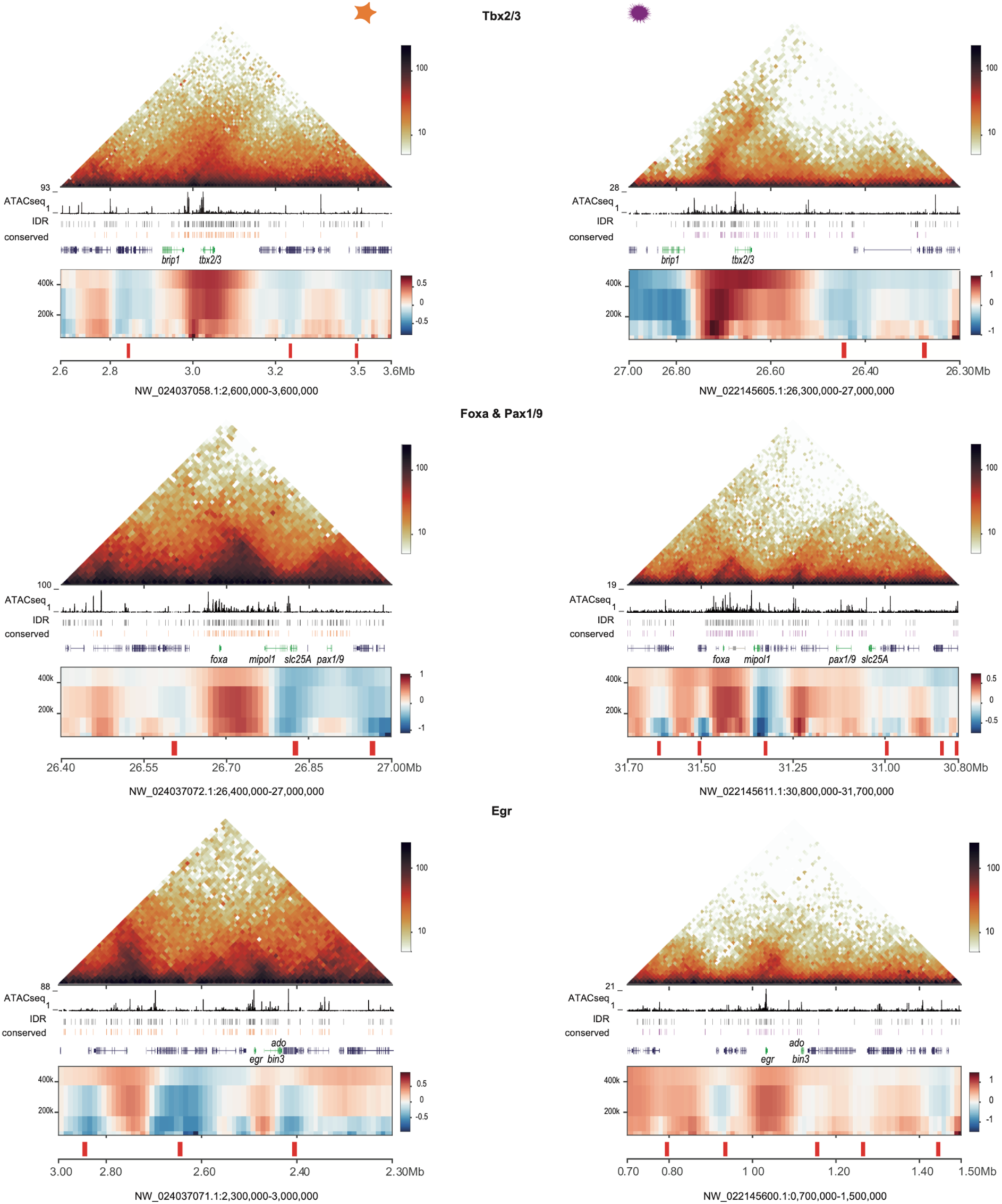
Examples of ancient conserved microsyntenic blocks located within TAD boundaries. Sea star (left panels) and sea urchin (right panels) genomic regions around four developmental genes included within ancient microsyntenic blocks (Tbx2/3, Foxa and Pax1/9, and Egr). From top to bottom: heatmaps showing normalized HiC signal at 10-kb resolution in late gastrula sea star and sea urchin embryos, ATAC-seq signal at late gastrula stage embryos of both species, IDR ATAC-seq peaks (pCREs), conserved ATAC-seq peaks (merging peaks conserved at the Valvatida+Asteroidea strata and Odontophora+Echinoidea strata), gene models (with those included within the conserved syntenic blocks colored in green), insulation scores and computationally called TAD boundaries. Note that the conserved block containing Foxa and Pax1/9 is split in two in the case of sea urchin.

**Extended Data Figure 3.**
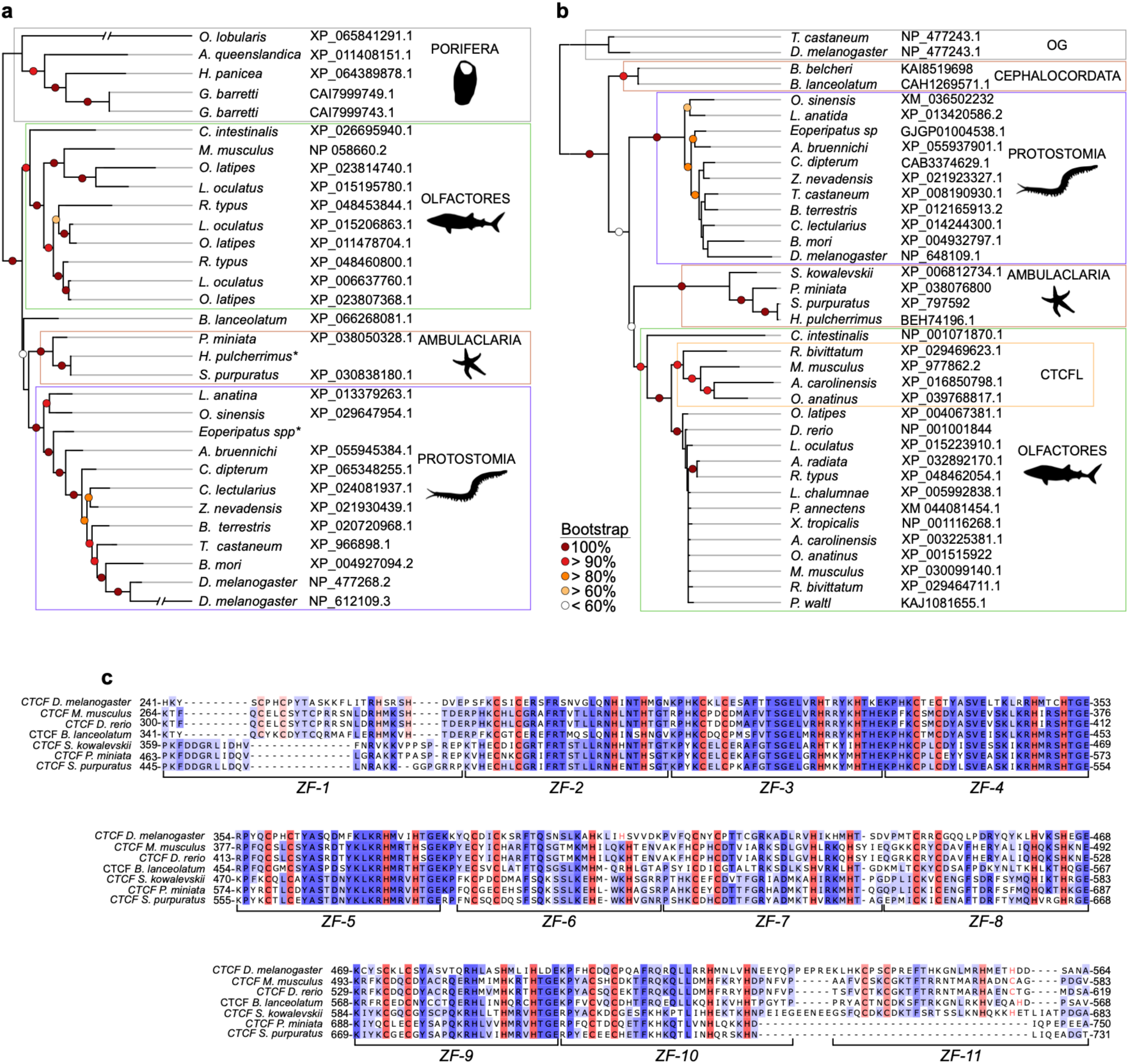
Cohesin and CTCF phylogeny and conservation. a, ML phylogenetic tree of cohesin subunit SA1-3 proteins. Sponge SA proteins were used as outgroups. b, ML phylogenetic tree of CTCF, including the tetrapod-specific paralog CTCFL (also known as BORIS), the outgroup (OG) branch includes the insect proteins CROOKED LEGS, a zinc finger containing family of transcription factors. c, Alignment of CTCF proteins from different bilaterians, showing the region containing its putatively ancestral 11 zinc finger domains, where ambulacrarian species have lost one or two domains.

**Extended Data Figure 4.**
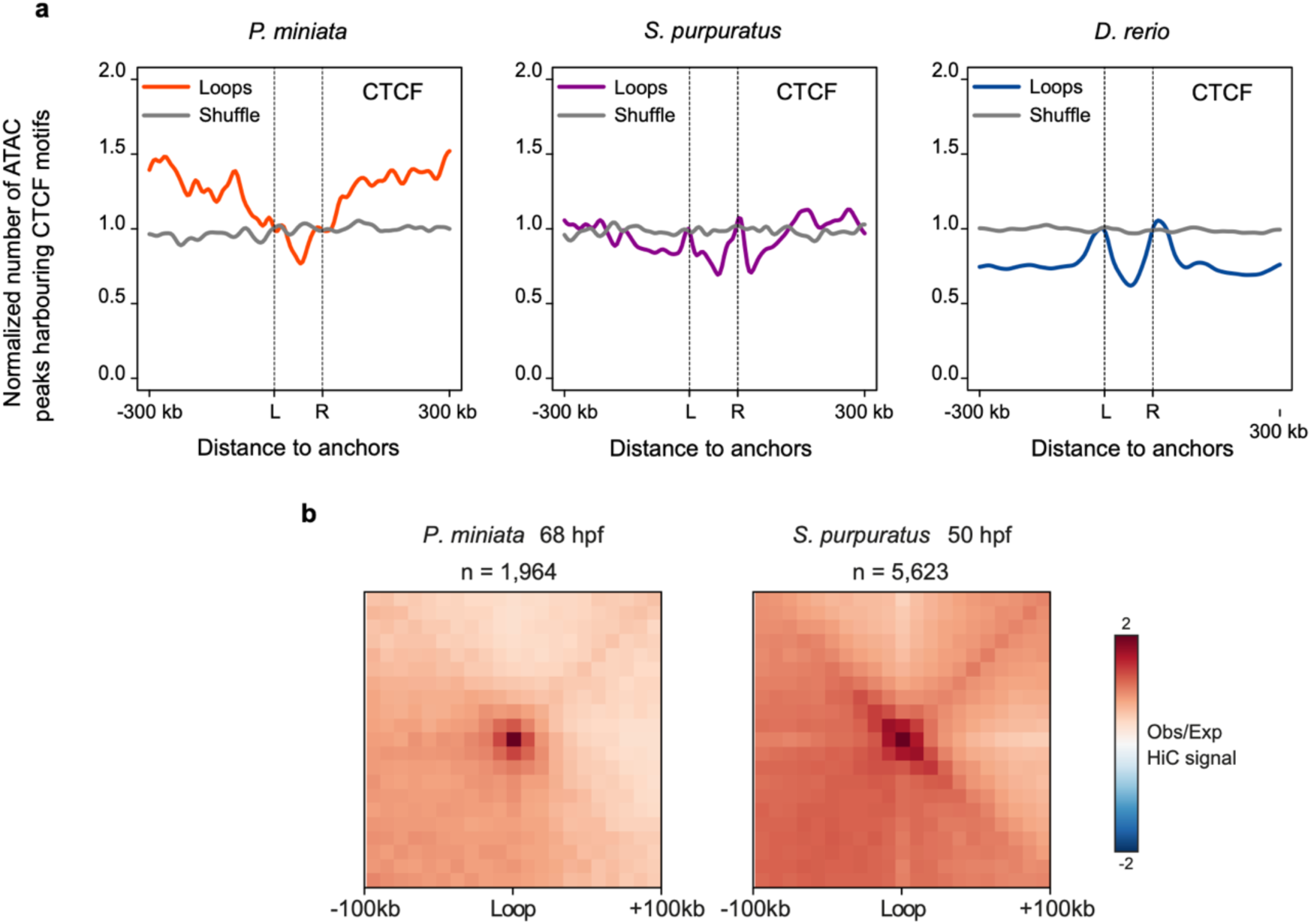
Chromatin loops in echinoderms. a, Normalized number of ATAC-seq peaks harboring CTCF motifs around HiC loops in *P. miniata* (left), *S. purpuratus* (middle) and *D. rerio* (right). Shuffle control is shown per each graph. b, Aggregate peak analyses of HiC loops in *P. miniata* (left) and *S. purpuratus* (right) late gastrula embryos. The observed vs. expected HiC signal is shown.

**Extended Data Fig. 5.**
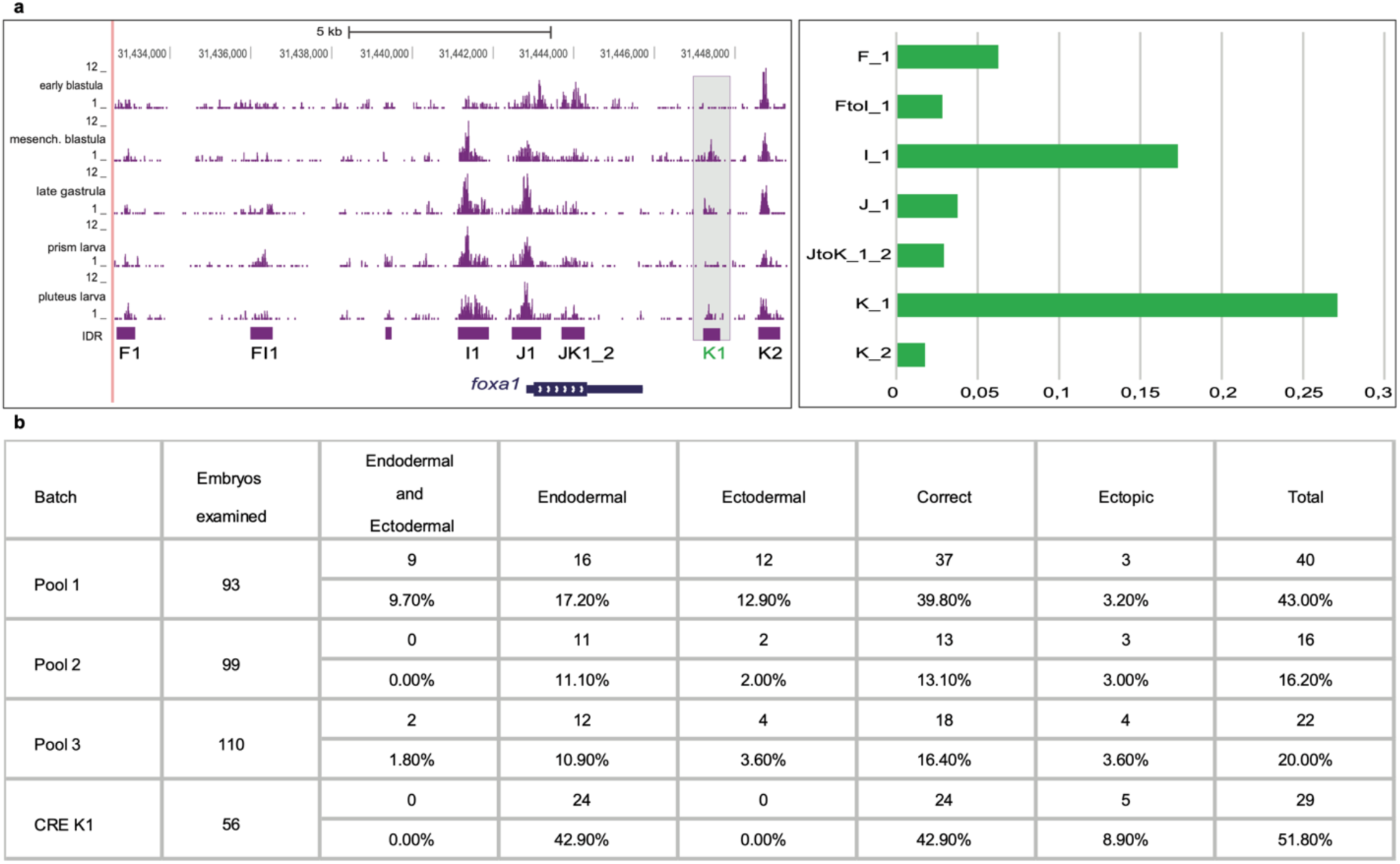
Functional assays of *foxa1* pCREs. a, Screenshot of sea urchin UCSC browser with ATAC-seq tracks around the *foxa1* (LOC110977664) gene and pCREs. Scoring table for embryos injected with *foxa1* CRE-Tag constructs at mesenchyme blastula stage. b, Relative GFP expression levels driven by *foxa1* CREs in sea urchin embryos at mesenchyme blastula stage. The embryos injected with the pool of *foxa1* CREs showing GFP expression in various domains were scored, highlighting the distribution of embryos with expression in both ectoderm and endoderm, only ectoderm or only endoderm, as well as the ones exhibiting GFP expression in regions ectopic to *foxa1* expression. This was also concordant with scoring done for the FIJ concatenate ^53^ since the majority of the expression is in the endoderm with oral ectoderm expression exhibited by fewer embryos.

**Extended Data Fig. 6.**
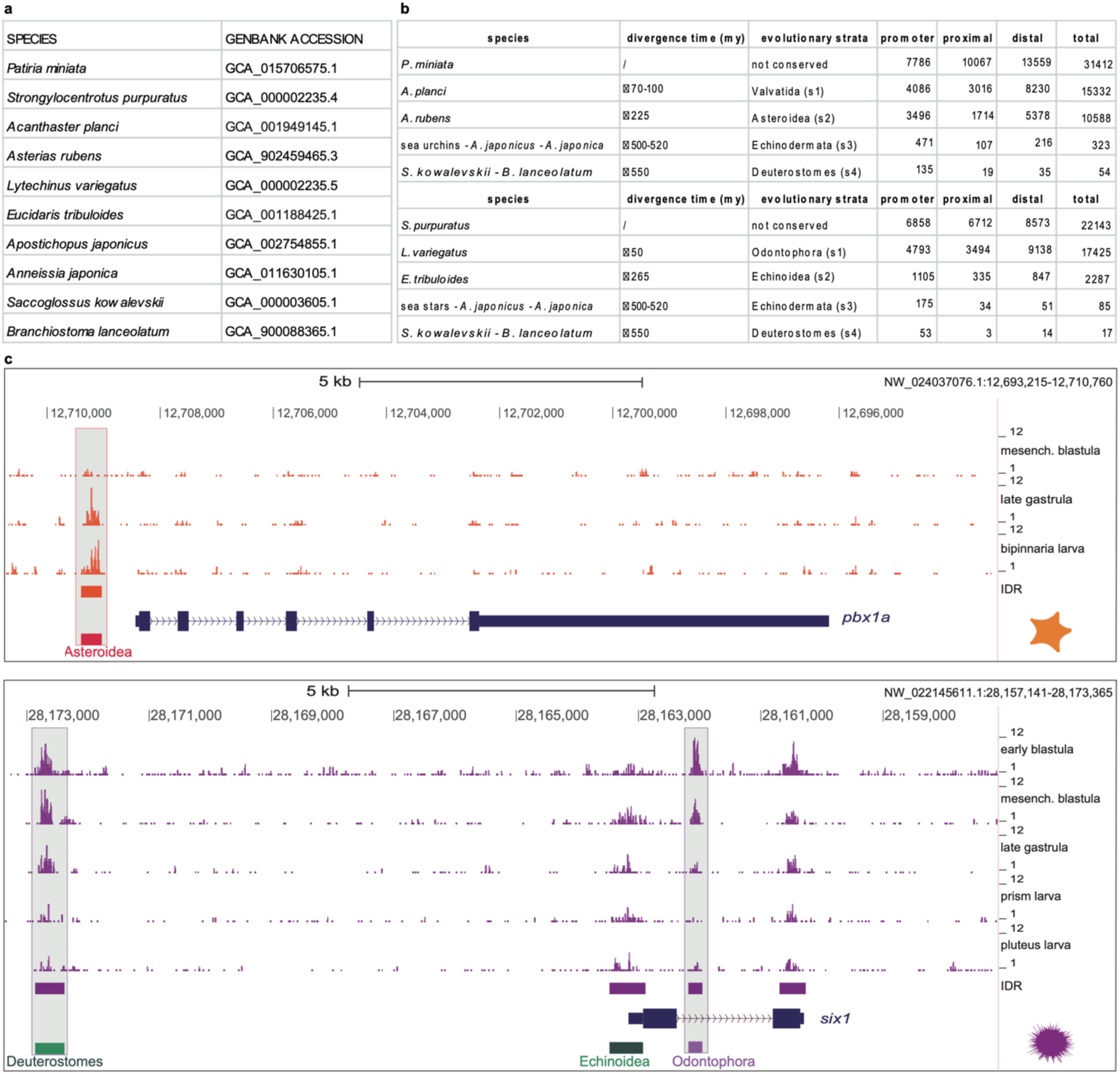
Evolutionary strata and developmental dynamics of sea urchin and sea star pCREs. **a**, Accession numbers of the genome assemblies included in the alignments. **b**, Counts of pCREs according to their evolutionary strata. **c**, Genome browser screenshots around the *P. miniata pbx1a* (LOC119720117, top panel) and the *S. purpuratus six1* (LOC110974175, lower panel) genes showing ATAC-seq tracks from different developmental stages (in orange and purple, respectively), IDR peaks (orange and purple bars) and gene models (dark blue). A conserved Asteroidea pCRE (red bar) with late developmental dynamics and Deuterostome and Odontophora pCREs (green bars) with early developmental dynamics are highlighted with gray boxes.

**Extended Data Fig. 7.**
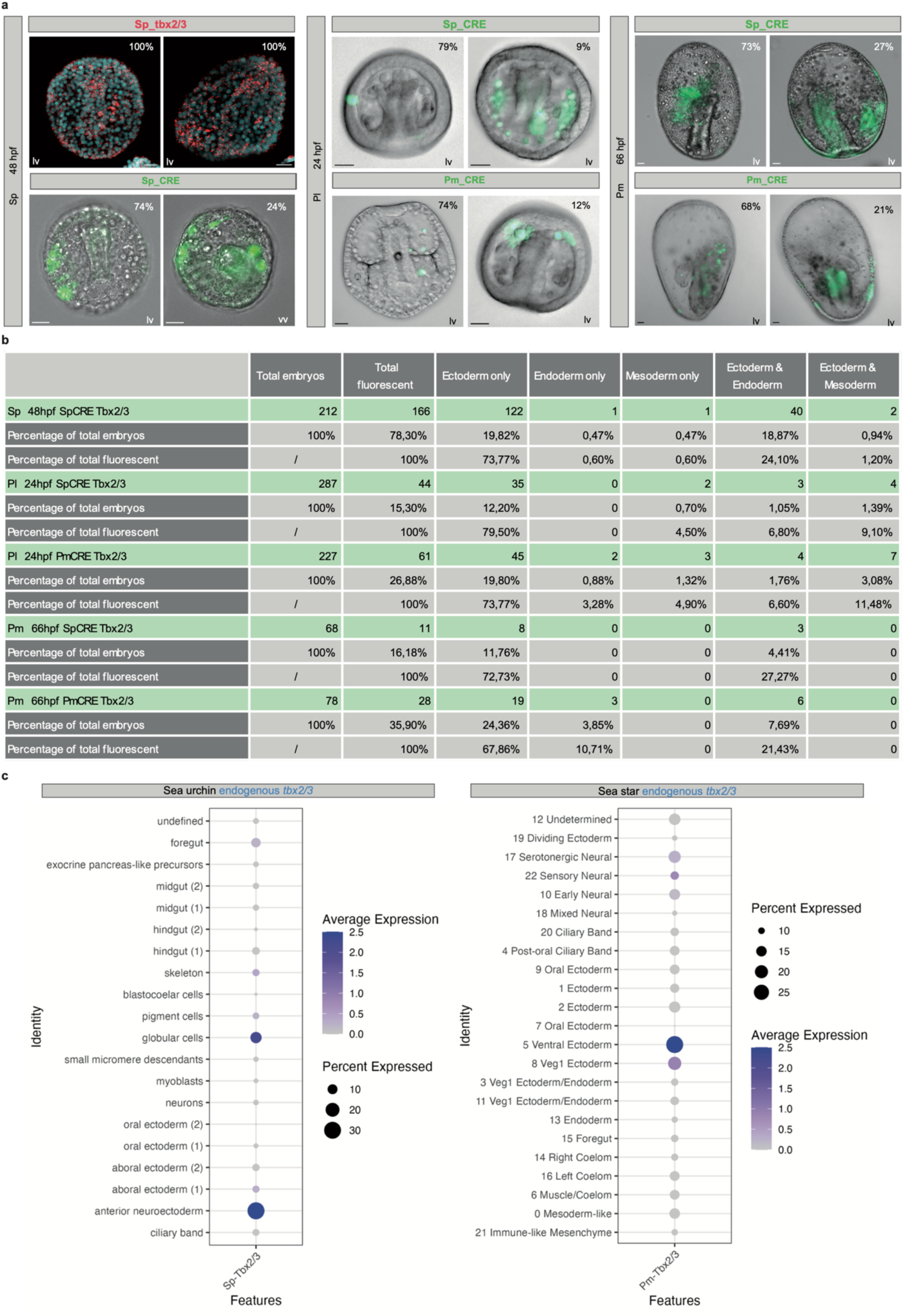
Functional assays of Tbx2/3 deeply conserved CRE. **a**, Numbers of embryos examined in functional assays with the two different CREs and percentages of expression in *S. purpuratus* (Sp), *P. lividus* (Pl), *P. miniata* (Pm). **b**, Expression pattern of *tbx2/3* by fluorescent in situ hybridization in *S. purpuratus* late gastrula (top left, focus on the ectoderm) and functional assays of *S. purpuratus* and *P. miniata* CREs in different species (*S. purpuratus* (Sp), *P. lividus* (Pl), *P. miniata* (Pm)) by GFP fluorescence. Scale bar 20 µm; lv, lateral view; vv, ventral view. **c**, The dot plots highlight the expression of endogenous *tbx2/3* as detected from scRNA-seq data analysis of gastrula stages in *S. purpuratus* ^49^, left column, and from snRNA-seq data analysis of gastrula stages in *P. miniata* ^73^, right column.

