## Extended Data Fig 1 for "Deep conservation of cis-regulatory elements and chromatin organization in echinoderms uncover ancestral regulatory features of animal genomes"

**a**

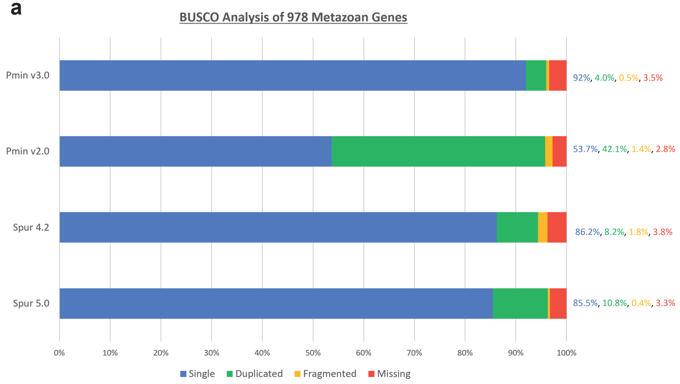


**b**

| Assembly | Size | Scaffolds | Scaffold N50 | Contigs | Contig N50 |
|---|---|---|---|---|---|
| Pmin v1.0 | 811 Mbp | 60,183 | 52.6 Kbp | 179,756 | 9.5 Kbp |
| Pmin v3.0 | 608 Mbp | 30 | 23.1 Mbp | 1,191 | 2.1 Mbp |
| Spur v4.2 | 991 Mbp | 31,896 | 420 Kbp | 146,29 | 16.8 Kbp |
| Spur v5.0 | 921 Mbp | 871 | 37 Mbp | 1,546 | 2.1 Mbp |

**c**

*P. miniata* v1.0

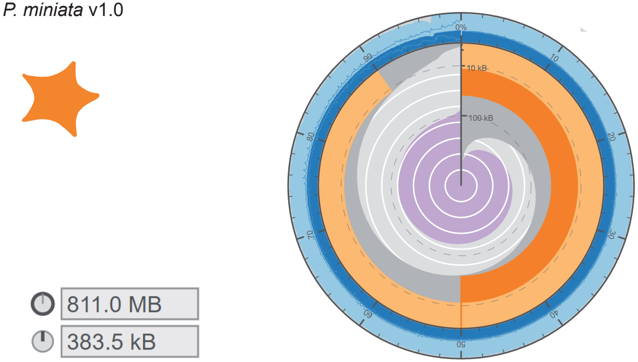


| Scaffold statistics | |
|---|---|
| Log 10 scaffold count | total 30 |
| Scaffold length | total 608 MB |
| Longest scaffold | 48.5 MB |
| N50 length | 23.1 MB |
| N90 length | 10.1 MB |
| **Assembly base composition** | |
| GC | 40.6% |
| AT | 59.3% |
| N | 0.1% |

*S. purpuratus* v4.2

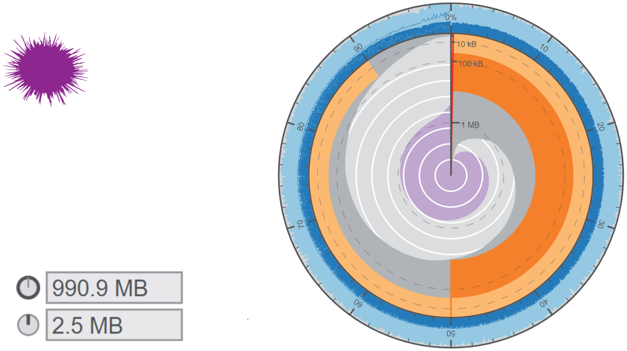


| Scaffold statistics | |
|---|---|
| Log 10 scaffold count | total 31,897 |
| Scaffold length | total 991 MB |
| Longest scaffold | 2.5 MB |
| N50 length | 419.6 KB |
| N90 length | 50.0 KB |
| **Assembly base composition** | |
| GC | 37.3% |
| AT | 53.8% |
| N | 8.9% |
