## Supplementary figures and images for "Deep conservation of cis-regulatory elements and chromatin organization in echinoderms uncover ancestral regulatory features of animal genomes"

### Extended Data Fig 2

#### Extended Data Fig. 2

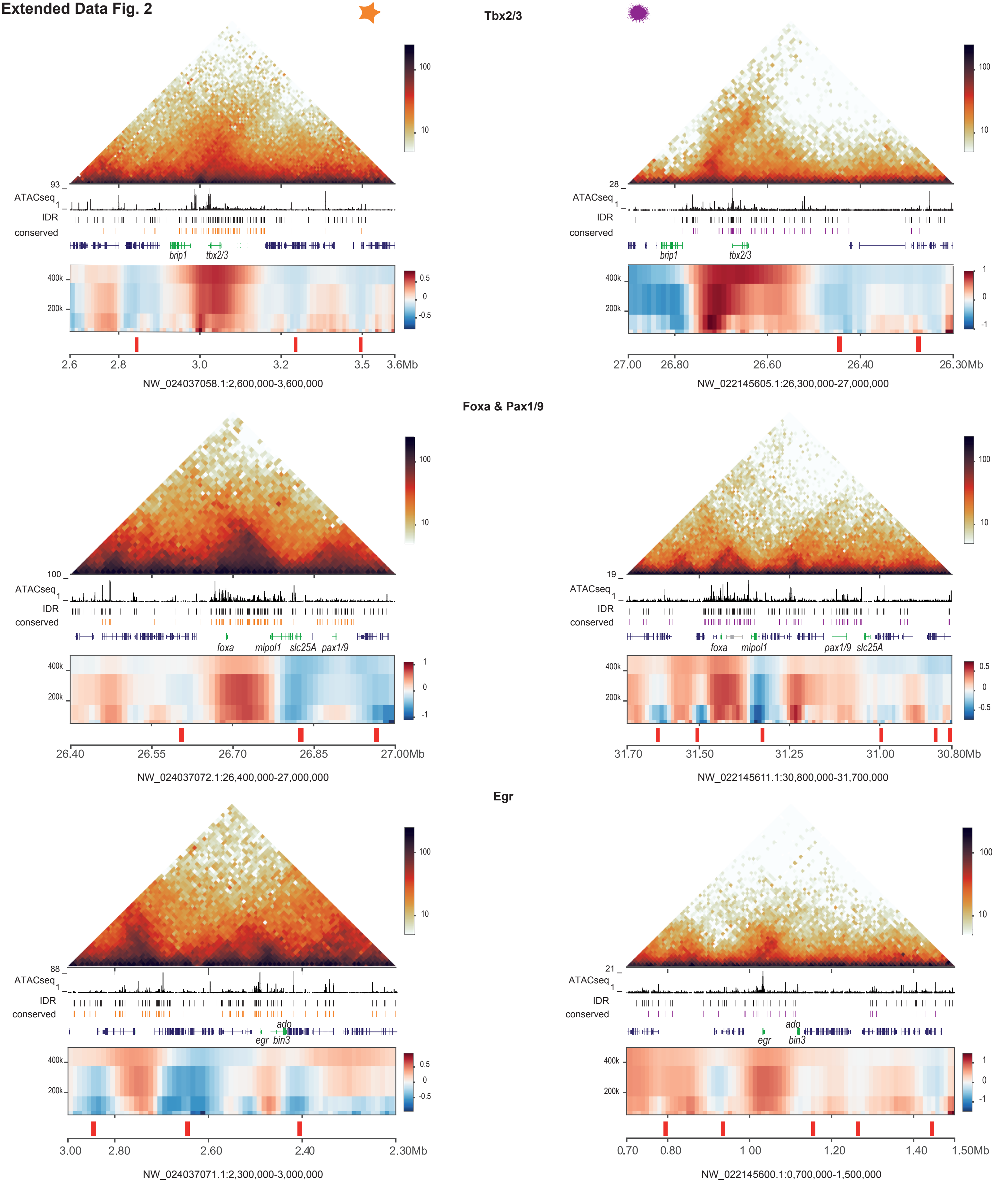

### Extended Data Fig 3

#### Extended Data Fig. 3

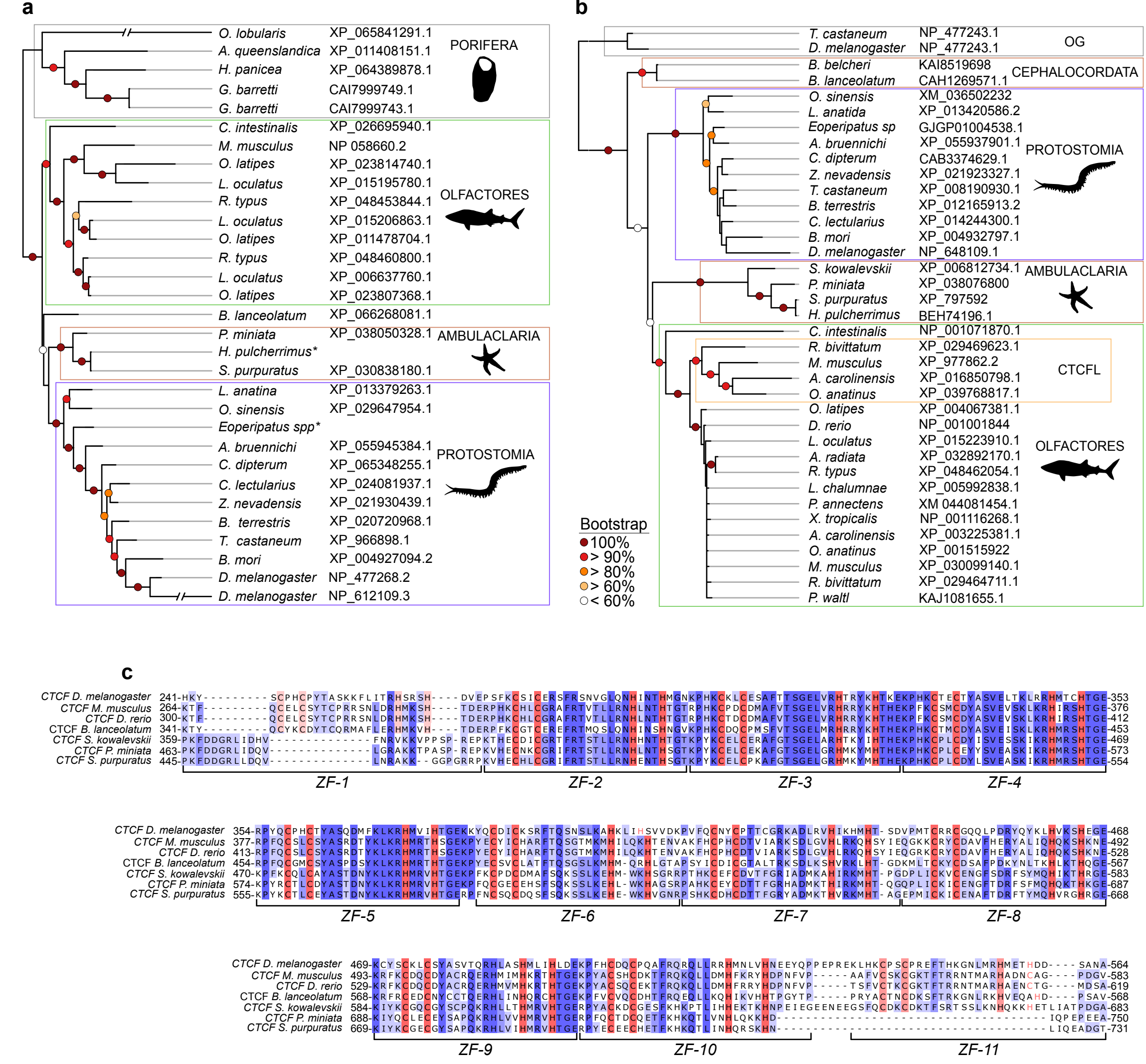

### Extended Data Fig 4

**Extended Data Fig. 4**

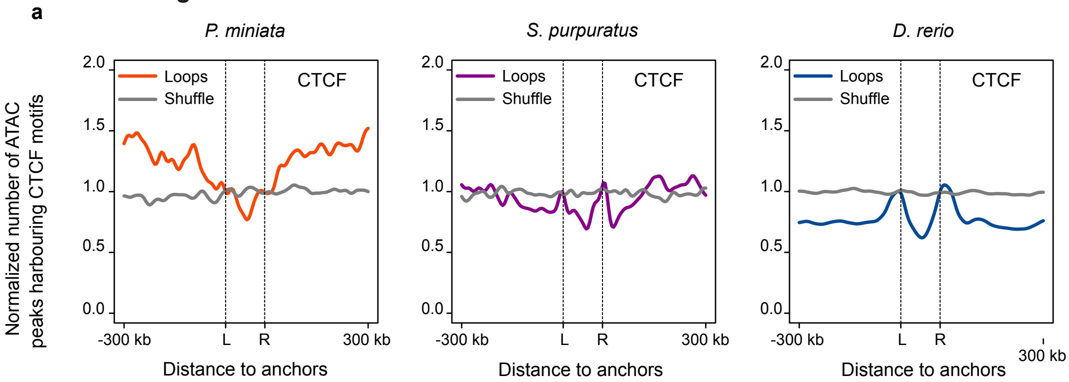


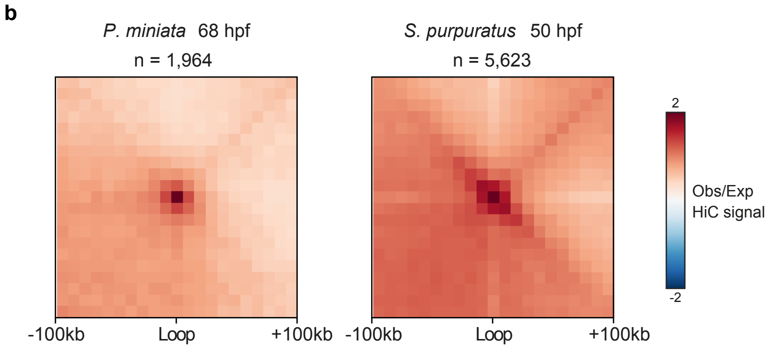
