## Extended Data Fig 5 for "Deep conservation of cis-regulatory elements and chromatin organization in echinoderms uncover ancestral regulatory features of animal genomes"

**a**

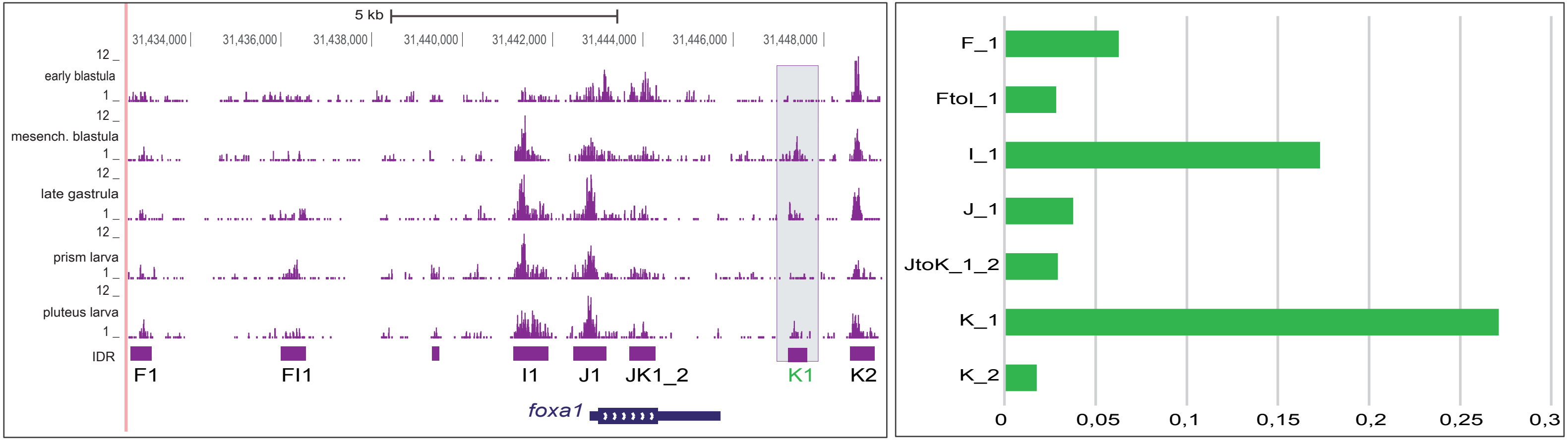


**b**

| Batch | Embryos examined | Endodermal and Ectodermal | Endodermal | Ectodermal | Correct | Ectopic | Total |
|---|---|---|---|---|---|---|---|
| Pool 1 | 93 | 9 | 16 | 12 | 37 | 3 | 40 |
| | | 9.70% | 17.20% | 12.90% | 39.80% | 3.20% | 43.00% |
| Pool 2 | 99 | 0 | 11 | 2 | 13 | 3 | 16 |
| | | 0.00% | 11.10% | 2.00% | 13.10% | 3.00% | 16.20% |
| Pool 3 | 110 | 2 | 12 | 4 | 18 | 4 | 22 |
| | | 1.80% | 10.90% | 3.60% | 16.40% | 3.60% | 20.00% |
| CRE K1 | 56 | 0 | 24 | 0 | 24 | 5 | 29 |
| | | 0.00% | 42.90% | 0.00% | 42.90% | 8.90% | 51.80% |
