## Extended Data Fig 6 for "Deep conservation of cis-regulatory elements and chromatin organization in echinoderms uncover ancestral regulatory features of animal genomes"

**a**

| SPECIES | GENBANK ACCESSION |
|---|---|
| *Patiria miniata* | GCA_015706575.1 |
| *Strongylocentrotus purpuratus* | GCA_000002235.4 |
| *Acanthaster planci* | GCA_001949145.1 |
| *Asterias rubens* | GCA_902459465.3 |
| *Lytechinus variegatus* | GCA_000002235.5 |
| *Eucidaris tribuloides* | GCA_001188425.1 |
| *Apostichopus japonicus* | GCA_002754855.1 |
| *Anneissia japonica* | GCA_011630105.1 |
| *Saccoglossus kowalevskii* | GCA_000003605.1 |
| *Branchiostoma lanceolatum* | GCA_900088365.1 |

**b**

| species | divergence time (my) | evolutionary strata | promoter | proximal | distal | total |
|---|---|---|---|---|---|---|
| *P. miniata* | / | not conserved | 7786 | 10067 | 13559 | 31412 |
| *A. planci* | 70-100 | Valvatida (s1) | 4086 | 3016 | 8230 | 15332 |
| *A. rubens* | 225 | Asteroidea (s2) | 3496 | 1714 | 5378 | 10588 |
| sea urchins - *A. japonicus* - *A. japonica* | 500-520 | Echinodermata (s3) | 471 | 107 | 216 | 323 |
| *S. kowalevskii* - *B. lanceolatum* | 550 | Deuterostomes (s4) | 135 | 19 | 35 | 54 |
| **species** | **divergence time (my)** | **evolutionary strata** | **promoter** | **proximal** | **distal** | **total** |
| *S. purpuratus* | / | not conserved | 6858 | 6712 | 8573 | 22143 |
| *L. variegatus* | 50 | Odontophora (s1) | 4793 | 3494 | 9138 | 17425 |
| *E. tribuloides* | 265 | Echinoidea (s2) | 1105 | 335 | 847 | 2287 |
| sea stars - *A. japonicus* - *A. japonica* | 500-520 | Echinodermata (s3) | 175 | 34 | 51 | 85 |
| *S. kowalevskii* - *B. lanceolatum* | 550 | Deuterostomes (s4) | 53 | 3 | 14 | 17 |

**c**

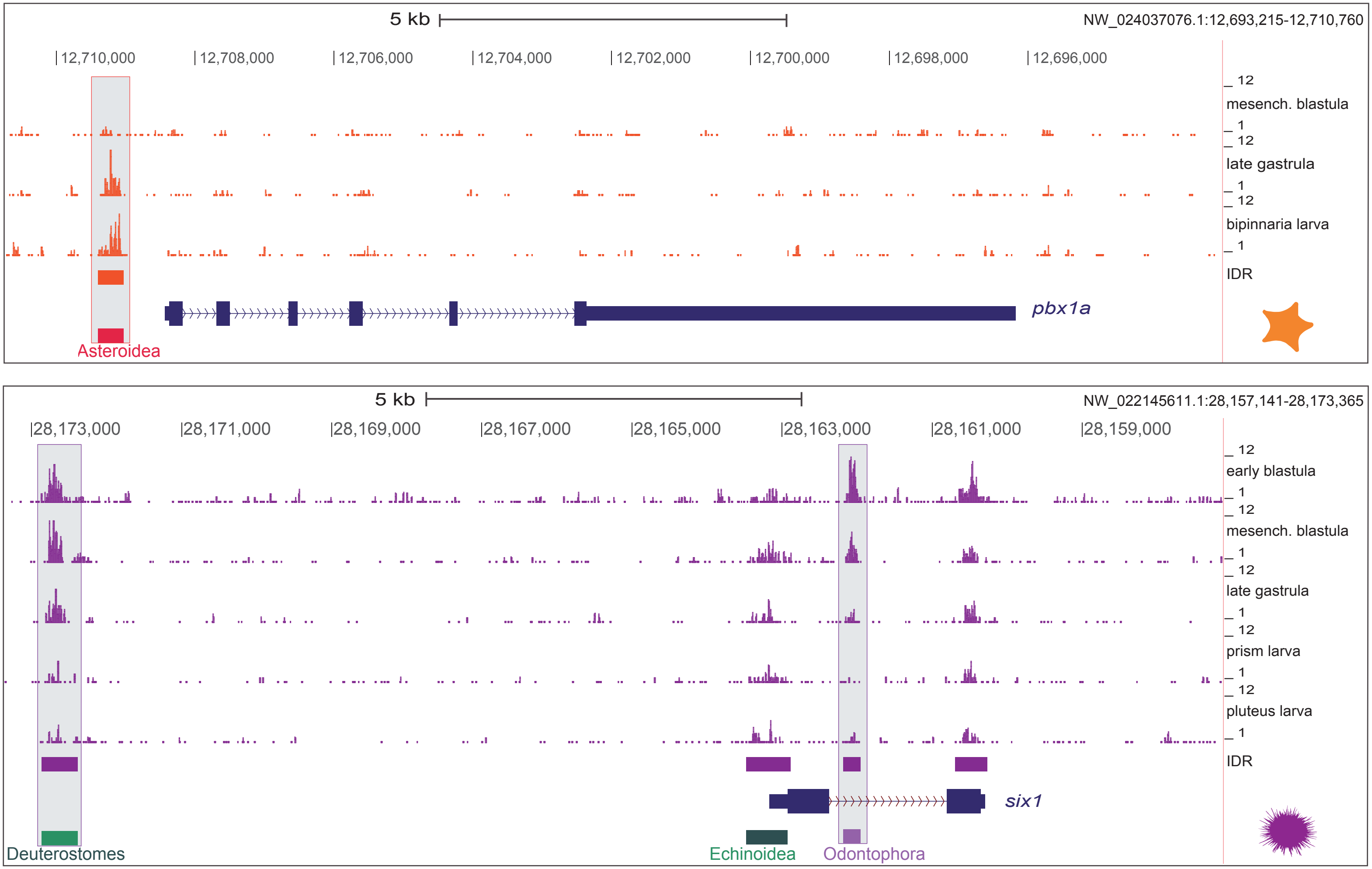
