## Extended Data Fig 7 for "Deep conservation of cis-regulatory elements and chromatin organization in echinoderms uncover ancestral regulatory features of animal genomes"

**a**

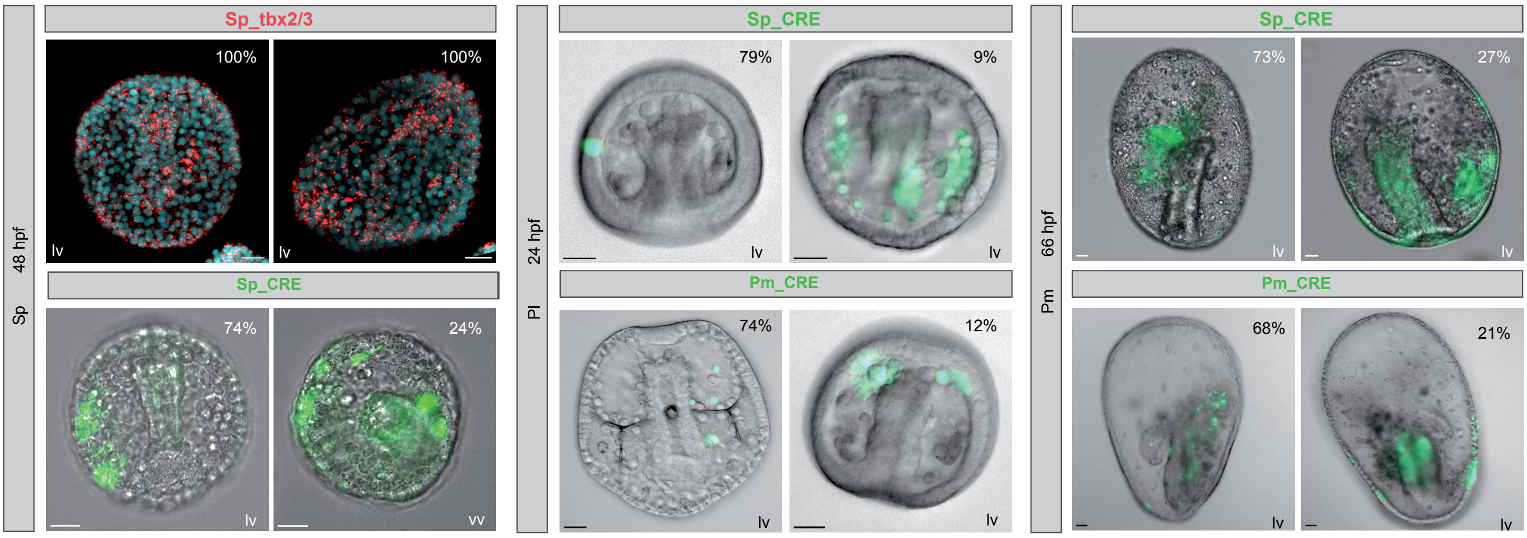


**b**

| | Total embryos | Total fluorescent | Ectoderm only | Endoderm only | Mesoderm only | Ectoderm & Endoderm | Ectoderm & Mesoderm |
|---|---|---|---|---|---|---|---|
| Sp 48hpf SpCRE Tbx2/3 | 212 | 166 | 122 | 1 | 1 | 40 | 2 |
| Percentage of total embryos | 100% | 78,30% | 19,82% | 0,47% | 0,47% | 18,87% | 0,94% |
| Percentage of total fluorescent | / | 100% | 73,77% | 0,60% | 0,60% | 24,10% | 1,20% |
| Pl 24hpf SpCRE Tbx2/3 | 287 | 44 | 35 | 0 | 2 | 3 | 4 |
| Percentage of total embryos | 100% | 15,30% | 12,20% | 0 | 0,70% | 1,05% | 1,39% |
| Percentage of total fluorescent | / | 100% | 79,50% | 0 | 4,50% | 6,80% | 9,10% |
| Pl 24hpf PmCRE Tbx2/3 | 227 | 61 | 45 | 2 | 3 | 4 | 7 |
| Percentage of total embryos | 100% | 26,88% | 19,80% | 0,88% | 1,32% | 1,76% | 3,08% |
| Percentage of total fluorescent | / | 100% | 73,77% | 3,28% | 4,90% | 6,60% | 11,48% |
| Pm 66hpf SpCRE Tbx2/3 | 68 | 11 | 8 | 0 | 0 | 3 | 0 |
| Percentage of total embryos | 100% | 16,18% | 11,76% | 0 | 0 | 4,41% | 0 |
| Percentage of total fluorescent | / | 100% | 72,73% | 0 | 0 | 27,27% | 0 |
| Pm 66hpf PmCRE Tbx2/3 | 78 | 28 | 19 | 3 | 0 | 6 | 0 |
| Percentage of total embryos | 100% | 35,90% | 24,36% | 3,85% | 0 | 7,69% | 0 |
| Percentage of total fluorescent | / | 100% | 67,86% | 10,71% | 0 | 21,43% | 0 |

**c**

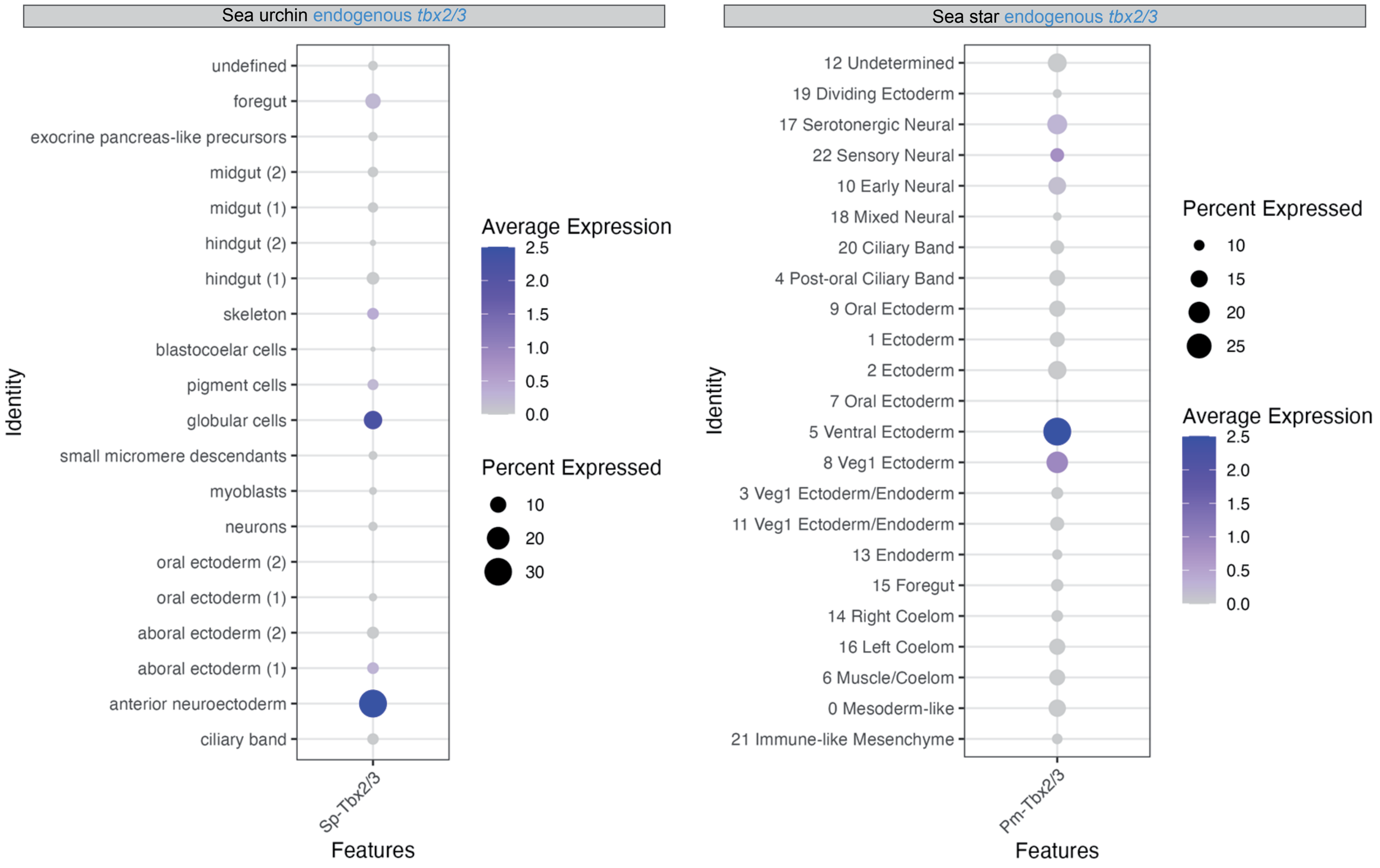
